# Novel deleterious nsSNPs within *MEFV* gene that could be used as Diagnostic Markers to Predict Hereditary Familial Mediterranean Fever: Using bioinformatics analysis

**DOI:** 10.1101/424796

**Authors:** Mujahed I. Mustafa, Tebyan A Abdelhameed, Fatima A. Abdelrhman, Soada Ahmed Osman, Mohamed A. Hassan

## Abstract

**Background:** Familial Mediterranean Fever (FMF) is the most common auto inflammatory disease (AID) affecting mainly the ethnic groups originating from Mediterranean basin, we aimed to identify the pathogenic SNPs in MEFV by computational analysis software.

**Methods:** We carried out in silico prediction of structural effect of each SNP using different bioinformatics tools to predict substitution influence on protein structure and function.

**Result:** 23 novel mutations out of 857 nsSNPs that are found to be deleterious effect on the MEFV structure and function.

**Conclusion:** This is the first in silico analysis in MEFV gene to prioritize SNPs for further genetic mapping studies. After using multiple bioinformatics tools to compare and rely on the results predicted, we found 23 novel mutations that may cause FMF disease and it could be used as diagnostic markers for Mediterranean basin populations.

## 1. Introduction

Familial Mediterranean fever is an autosomal recessive inherited inflammatory disease (1, 9, 12) (however, it has been observed that a substantial number of patients with clinical FMF possess only one demonstrable *MEFV* mutation (2, 3)) that is principally seen in different countries (4, 5, 6, 7, 8). However, patients from different ethnicities (such as Japan) are being increasingly recognized (9, 10), the carrier frequency for *MEFV* genetic variants in the population in the Mediterranean basin to be about 8% (11). Most cases of FMF usually present with acute abdominal pain and fever (1, 5, 12), both of which are also the main causes of referral in the emergency department.(13) all these factors may help For medical treatment, colchicine is the first line therapy(14), but in resistant cases (<10% of patients) (15) or affects the responsiveness to Colchicine(16) other anti-inflammatory drugs can be used for extra anti-inflammatory effect(17). If FMF not be treated, may be an etiologic factor for colonic LNH in children.(18) *MEFV* gene is localized on 16p13.3 of chromosome 16 at position 13.3 which consist of 10 exons with 21600 bp (12, 19). The disease is characterized by recurrent febrile episodes and inflammation in the form of sterile polyserositis Amyloid protein involved in inflammatory amyloidosis was named AA (amyloid‐associated) protein and its circulating precursor was named SAA (serum amyloid‐associated). Amyloidosis of the AA type is the most severe complication of the disease. The gene responsible for FMF, *MEFV*, encodes a protein called (pyrin or marenostrin) and is expressed mainly in neutrophils. (12, 19)

The definition of the *MEFV* gene has permitted genetic diagnosis of the disease. Nevertheless, as studies have unwrapped molecular data, problems have arisen with the clinical definitions of the disease.(20) FMF is caused by mutations in the *MEFV* missense SNPs (we were focusing on SNPs which are located in the coding region because it is much important in disease causing potential, which are responsible for amino acid residue substitutions resulting in functional diversity of proteins in humans) (20) coding for pyrin, which is a component of inflammasome functioning in inflammatory response and production of interleukin-1β (IL-1β). Recent studies have shown that pyrin recognizes bacterial modifications in Rho GTPases, which results in inflammasome activation and increase in IL-1β. Pyrin does not directly recognize Rho modification but probably affected by Rho effector kinase, which is a downstream event in the actin cytoskeleton pathway.(19, 21, 22)

The aim of this study was to identify the pathogenic SNPs in *MEFV* using insilico prediction software, and to determine the structure, function and regulation of their respective proteins. This is the first in silico analysis in *MEFV* gene to prioritize SNPs for further genetic mapping studies. The usage of insilico approach has strong impact on the identification of candidate SNPs since they are easy and less costly, and can facilitate future genetic studies (23).

## 2. Method

### 2.1 Data mining

The data on human *MEFV* gene was collected from National Center for Biological Information (NCBI) web site (24). The SNP information (protein accession number and SNP ID) of the *MEFV* gene was retrieved from the NCBI dbSNP (http://www.ncbi.nlm.nih.gov/snp/) and the protein sequence was collected from Swiss Prot databases (http://expasy.org/). (25)

### 2.2 SIFT

SIFT is a sequence homology-based tool (26) that sorts intolerant from tolerant amino acid substitutions and predicts whether an amino acid substitution in a protein will have a phenotypic Effect. Considers the position at which the change occurred and the type of amino acid change. Given a protein sequence, SIFT chooses related proteins and obtains an alignment of these proteins with the query. Based on the amino acids appearing at each position in the alignment, SIFT calculates the probability that an amino acid at a position is tolerated conditional on the most frequent amino acid being tolerated. If this normalized value is less than a cutoff, the substitution is predicted to be deleterious. SIFT scores <0.05 are predicted by the algorithm to be intolerant or deleterious amino acid substitutions, whereas scores >0.05 are considered tolerant. It is available at (http://sift.bii.a-star.edu.sg/).

### 2.3 Polyphen-2

It is a software tool (27) to predict possible impact of an amino acid substitution on both structure and function of a human protein by analysis of multiple sequence alignment and protein 3D structure, in addition it calculates position-specific independent count scores (PSIC) for each of two variants, and then calculates the PSIC scores difference between two variants. The higher a PSIC score difference, the higher the functional impact a particular amino acid substitution is likely to have. Prediction outcomes could be classified as probably damaging, possibly damaging or benign according to the value of PSIC as it ranges from (0_1); values closer to zero considered benign while values closer to 1 considered probably damaging and also it can be indicated by a vertical black marker inside a color gradient bar, where green is benign and red is damaging. nsSNPs that predicted to be intolerant by Sift has been submitted to Polyphen as protein sequence in FASTA format that obtained from UniproktB /Expasy after submitting the relevant ensemble protein (ESNP) there, and then we entered position of mutation, native amino acid and the new substituent for both structural and functional predictions. PolyPhen version 2.2.2 is available at http://genetics.bwh.harvard.edu/pph2/index.shtml

### 2.4 Provean

Provean is a software tool (28) which predicts whether an amino acid substitution or indel has an impact on the biological function of a protein. it is useful for filtering sequence variants to identify nonsynonymous or indel variants that are predicted to be functionally important. It is available at (https://rostlab.org/services/snap2web/).

### 2.5 SNAP2

Functional effects of mutations are predicted with SNAP2 (29). SNAP2 is a trained classifier that is based on a machine learning device called “neural network”. It distinguishes between effect and neutral variants/non-synonymous SNPs by taking a variety of sequence and variant features into account. The most important input signal for the prediction is the evolutionary information taken from an automatically generated multiple sequence alignment. Also structural features such as predicted secondary structure and solvent accessibility are considered. If available also annotation (i.e. known functional residues, pattern, regions) of the sequence or close homologs are pulled in. In a cross-validation over 100,000 experimentally annotated variants, SNAP2 reached sustained two-state accuracy (effect/neutral) of 82% (at an AUC of 0.9). In our hands this constitutes an important and significant improvement over other methods. It is available at (https://rostlab.org/services/snap2web/).

### 2.6 PHD-SNP

An online Support Vector Machine (SVM) based classifier, is optimized to predict if a given single point protein mutation can be classified as disease-related or as a neutral polymorphism. it is available at: (http://snps.biofold.org/phd-snp/phdsnp.html)

### 2.7 SNP& Go

SNPs&GO is an algorithm developed in the Laboratory of Biocomputing at the University of Bologna directed by Prof. Rita Casadio. SNPs&GO is an accurate method that, starting from a protein sequence, can predict whether a variation is disease related or not by exploiting the corresponding protein functional annotation. SNPs&GO collects in unique framework information derived from protein sequence, evolutionary information, and function as encoded in the Gene Ontology terms, and outperforms other available predictive methods.(30) It is available at (http://snps.biofold.org/snps-and-go/snps-and-go.html)

### 2.8 P-Mut

PMUT a web-based tool (31) for the annotation of pathological variants on proteins, allows the fast and accurate prediction (approximately 80% success rate in humans) of the pathological character of single point amino acidic mutations based on the use of neural networks. It is available at (http://mmb.irbbarcelona.org/PMut).

### 2.9 I-Mutant 3.0

I-Mutant 3.0 Is a neural network based tool(32) for the routine analysis of protein stability and alterations by taking into account the single-site mutations. The FASTA sequence of protein retrieved from UniProt is used as an input to predict the mutational effect on protein stability. It is available at (http://gpcr2.biocomp.unibo.it/cgi/predictors/I-Mutant3.0/I-Mutant3.0.cgi).

### 2.10 BioEdit

BioEdit is an easy-to-use biological sequence alignment editor. This free software is intended to supply a single program that can handle most simple sequence and alignment editing and manipulation functions as well as a few basic sequences analyses. It is available for download at (http://www.mbio.ncsu.edu/bioedit/bioedit.html)

### 2.11 Modeling nsSNP locations on protein structure

Project hope is a new online web-server to search protein 3D structures (if available) by collecting structural information from a series of sources, including calculations on the 3D coordinates of the protein, sequence annotations from the UniProt database, and predictions by DAS services. Protein sequences were submitted to project hope server in order to analyze the structural and conformational variations that have resulted from single amino acid substitution corresponding to single nucleotide substitution. It is available at (http://www.cmbi.ru.nl/hope)

### 2.12 GeneMANIA

We submitted genes and selected from a list of data sets that they wish to query. GeneMANIA (33) approach to know protein function prediction integrate multiple genomics and proteomics data sources to make inferences about the function of unknown proteins. It is available at (http://www.genemania.org/)

## 3. Results and Discussion

### Result

**Table (1):** Damaging or Deleterious or effect nsSNPs associated variations predicted by various softwares:

| Amino Acid Change | SIFT |  | Polyphen |  | PROVEAN |  | SNAP2 |  |
| --- | --- | --- | --- | --- | --- | --- | --- | --- |
|  | prediction | Score | Prediction | score | Score | Prediction (cutoff= - 2.5) | prediction | score |
| S749Y | DAMAGING | 0 | PROBABLY DAMAGING | 1.000 | -3.116 | Deleterious | effect | 64 |
| F743S | DAMAGING | 0 | PROBABLY DAMAGING | 1.000 | -5.563 | Deleterious | effect | 68 |
| Y741C | DAMAGING | 0 | PROBABLY DAMAGING | 1.000 | -6.035 | Deleterious | effect | 77 |
| F731V | DAMAGING | 0 | PROBABLY DAMAGING | 1.000 | -5.159 | Deleterious | effect | 81 |
| I720T | DAMAGING | 0 | PROBABLY DAMAGING | 1.000 | -3.639 | Deleterious | effect | 58 |
| L709R | DAMAGING | 0 | PROBABLY DAMAGING | 1.000 | -4.311 | Deleterious | effect | 77 |
| V691G | DAMAGING | 0 | PROBABLY DAMAGING | 1.000 | -4.667 | Deleterious | effect | 66 |
| W689R | DAMAGING | 0 | PROBABLY DAMAGING | 1.000 | -10.132 | Deleterious | effect | 89 |
| G668R | DAMAGING | 0 | PROBABLY DAMAGING | 1.000 | -6.287 | Deleterious | effect | 92 |
| V659F | DAMAGING | 0 | PROBABLY DAMAGING | 1.000 | -3.811 | Deleterious | effect | 64 |
| F636C | DAMAGING | 0 | PROBABLY DAMAGING | 1.000 | -6.49 | Deleterious | effect | 79 |
| R461W | DAMAGING | 0 | PROBABLY DAMAGING | 1.000 | -5.456 | Deleterious | effect | 68 |
| H407Q | DAMAGING | 0 | PROBABLY DAMAGING | 1.000 | -7.335 | Deleterious | effect | 41 |
| H407R | DAMAGING | 0 | PROBABLY DAMAGING | 1.000 | -7.332 | Deleterious | effect | 51 |
| H404R | DAMAGING | 0 | PROBABLY<br>DAMAGING | 1.000 | -7.349 | Deleterious | effect | 75 |
| C398Y | DAMAGING | 0 | PROBABLY<br>DAMAGING | 1.000 | -10.314 | Deleterious | effect | 51 |
| C395Y | DAMAGING | 0 | PROBABLY<br>DAMAGING | 1.000 | -10.262 | Deleterious | effect | 19 |
| C395F | DAMAGING | 0 | PROBABLY<br>DAMAGING | 1.000 | -10.315 | Deleterious | effect | 27 |
| C395R | DAMAGING | 0 | PROBABLY<br>DAMAGING | 1.000 | -11.074 | Deleterious | effect | 27 |
| H378Q | DAMAGING | 0 | PROBABLY<br>DAMAGING | 1.000 | -5.886 | Deleterious | effect | 38 |
| H378Y | DAMAGING | 0 | PROBABLY<br>DAMAGING | 1.000 | -4.884 | Deleterious | effect | 45 |
| C375R | DAMAGING | 0 | PROBABLY<br>DAMAGING | 1.000 | -8.429 | Deleterious | effect | 66 |
| L86P | DAMAGING | 0 | PROBABLY<br>DAMAGING | 1.000 | -4.1 | Deleterious | effect | 19 |

## DISSCUSSION

23 novel mutations have been found *(see table 3) that* effect on the stability and function of the *MEFV* gene using bioinformatics tools. The methods used were based on different aspects and parameters describing the pathogenicity and provide clues on the molecular level about the effect of mutations. It was not easy to predict the pathogenic effect of SNPs using single method. Therefore, multiple methods were used to compare and rely on the results predicted. In this study we used different in silico prediction algorithms: SIFT, PolyPhen-2, Provean, SNAP2, SNP&GO, PHD-SNP, P-MUT and I-Mutant 3.0. (see figure 1).

**Figure 1:**
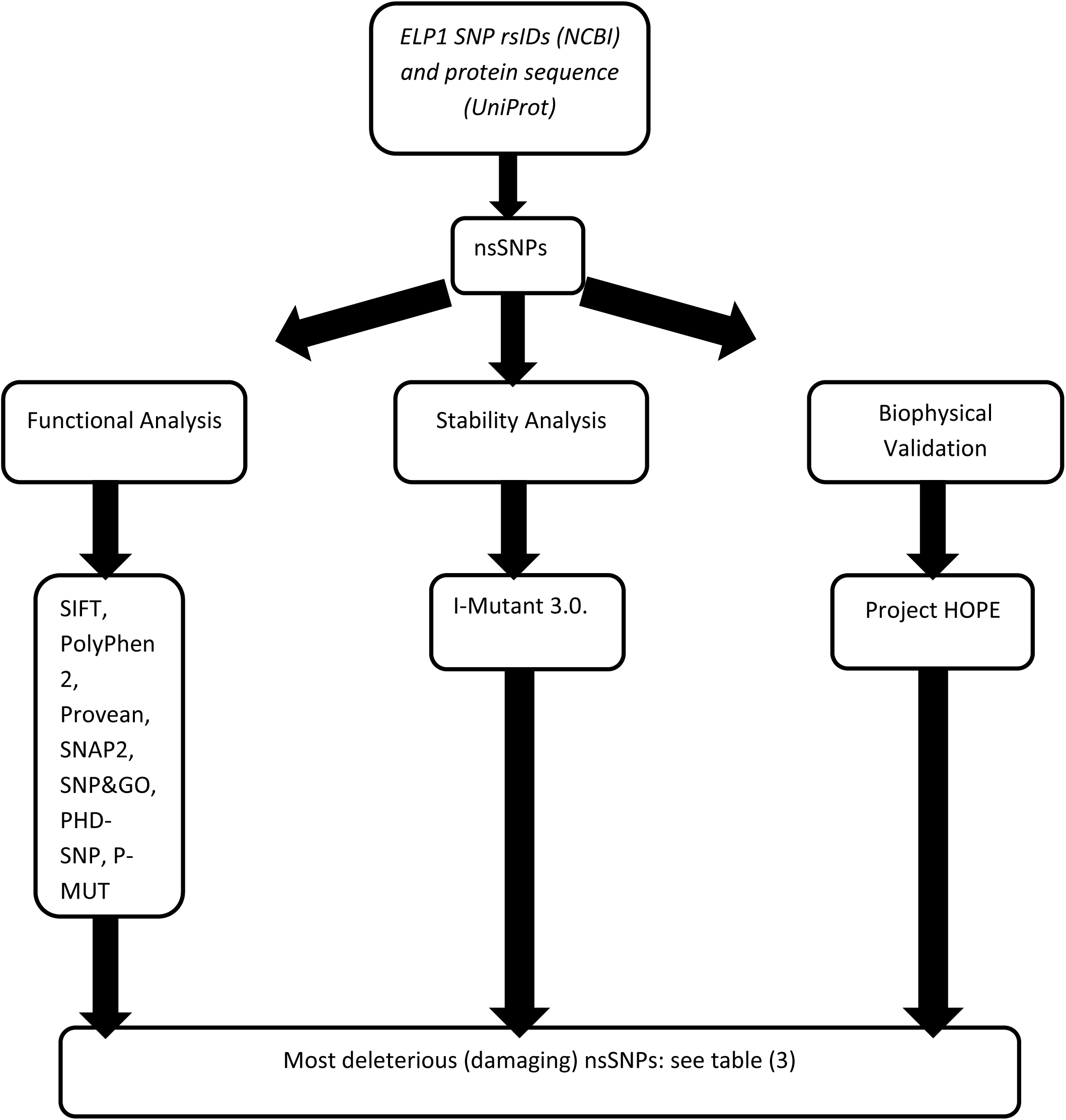
Diagrammatic representation of *IKBKAP* gene in silico work flow.

**Figure 2:**
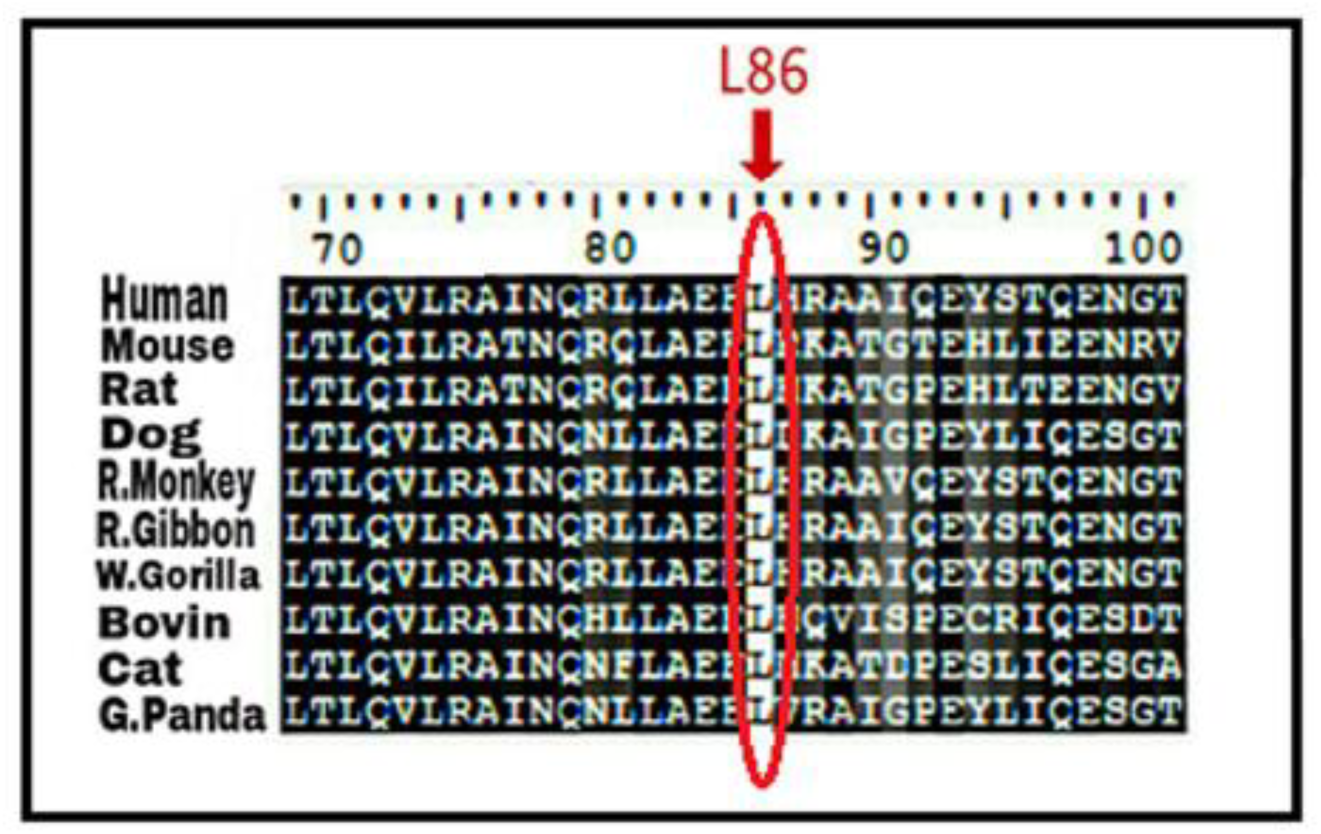
Alignments of 10 amino acid sequences of *MEFV* demonstrating that the residues predicted to be mutated in our band (indicated by red arrow) are evolutionarily conserved across species. Sequences Alignment were done by BioEdit (v7.2.5).

This study identified the total number of nsSNP in Homo sapiens located in coding region of *MEFV* gene, were investigate in dbSNP/NCBI Database (24) out of 2369 there are 856 nsSNPs (missense mutations), were submitted to SIFT server, PolyPhen-2 server, Provean sever and SNAP2 respectively, 392 SNPs were predicted to be deleterious in SIFT server. In PolyPhen-2 server, the result showed that 453 were found to be damaging (147 possibly damaging and 306 probably damaging showed deleterious). In Provean server our result showed that 244 SNPs were predicted to be deleterious. While in SNAP2 server the result showed that 566 SNPs were predicted to be Effect. The differences in prediction capabilities refer to the fact that every prediction algorithm uses different sets of sequences and alignments. In table (2) we were submitted four positive results from SIFT, PolyPhen-2, Provean and SNAP2 to observe the disease causing one by SNP&GO, PHD-SNP and P-Mut servers.

**Table (2):**
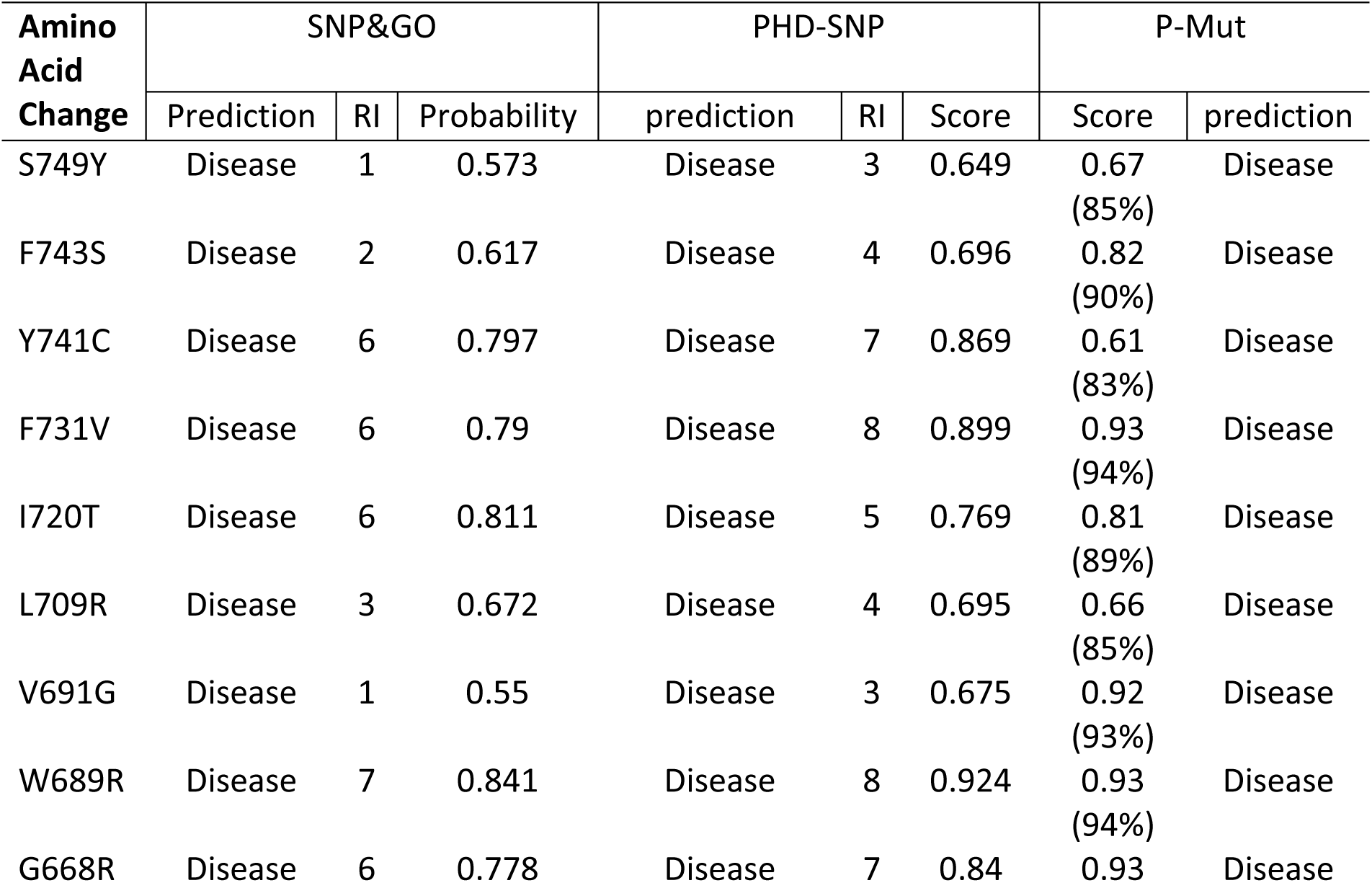
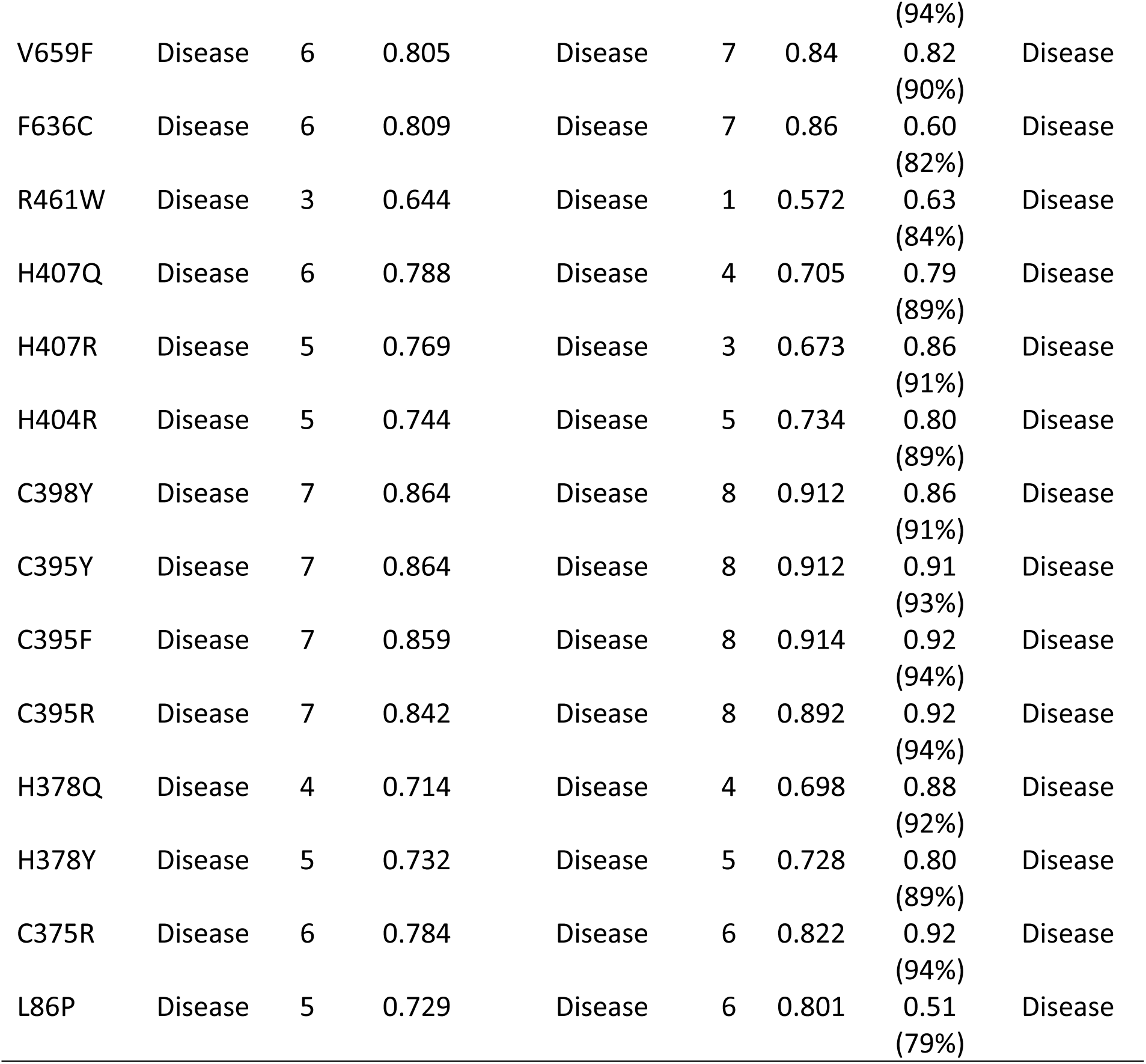
Disease effect nsSNPs associated variations predicted by various softwares:

In SNP&GO, PHD-SNP and P-Mut softwares were used to predict the association of SNPs with disease. According to SNP&GO, PHD-SNP and P-Mut (70, 91 and 58 SNPs respectively) were found to be disease related SNPs. We selected the triple disease related SNPs only in 3 softwares for further analysis by I-Mutant 3.0, Table (3). While I-Mutant result revealed that the protein stability decreased which destabilize the amino acid interaction (S749Y, F743S, Y741C, F731V, I720T, L709R, V691G, W689R, G668R, V659F, F636C, H407Q, H407R, H404R, C398Y, H378Q, H378Y, and L86P). While (C375R, C395F, C395R, C395Y, R461W) were found to be increased the protein stability (Table 3).

**Table (3):** stability analysis predicted by I-Mutant version 3.0 (also Show the 23 novel mutations):

| Amino Acid Change | SVM2 Prediction Effect | RI | DDG Value Prediction |
| --- | --- | --- | --- |
| S749Y | Decrease | 0 | -0.2 |
| F743S | Decrease | 6 | -1.16 |
| Y741C | Decrease | 8 | -2.5 |
| F731V | Decrease | 6 | -1.52 |
| I720T | Decrease | 4 | -0.92 |
| L709R | Decrease | 3 | -0.56 |
| V691G | Decrease | 7 | -1.25 |
| W689R | Decrease | 7 | -0.73 |
| G668R | Decrease | 4 | -0.37 |
| V659F | Decrease | 2 | -0.19 |
| F636C | Decrease | 8 | -1.28 |
| R461W | Increase | 1 | -0.01 |
| H407Q | Decrease | 8 | -1.48 |
| H407R | Decrease | 5 | -1.1 |
| H404R | Decrease | 1 | -0.06 |
| C398Y | Decrease | 2 | -0.09 |
| C395Y | Increase | 4 | 0.26 |
| C395F | Increase | 1 | 0.04 |
| C395R | Increase | 4 | 0.13 |
| H378Q | Decrease | 4 | -0.68 |
| H378Y | Decrease | 3 | -0.26 |
| C375R | Increase | 2 | -0.01 |
| L86P | Decrease | 2 | -0.56 |

Project HOPE software was used to submit the 23 most deleterious and damaging nsSNPs (figures from 3 to 25), L86P: Proline (The mutant residue) is smaller than Leucine (the wild-type residue) this might lead to loss of interactions. The wild-type and mutant amino acids differ in size. The mutation is located within a domain, annotated in UniProt as Pyrin. The mutation introduces an amino acid with different properties, which can disturb this domain and abolish its function. The wild-type residue is located in a region annotated in UniProt to form an α-helix. Proline disrupts an α-helix when not located at one of the first 3 positions of that helix. In case of the mutation at hand, the helix will be disturbed and this can have severe effects on the structure of the protein.

**Figure 3:**
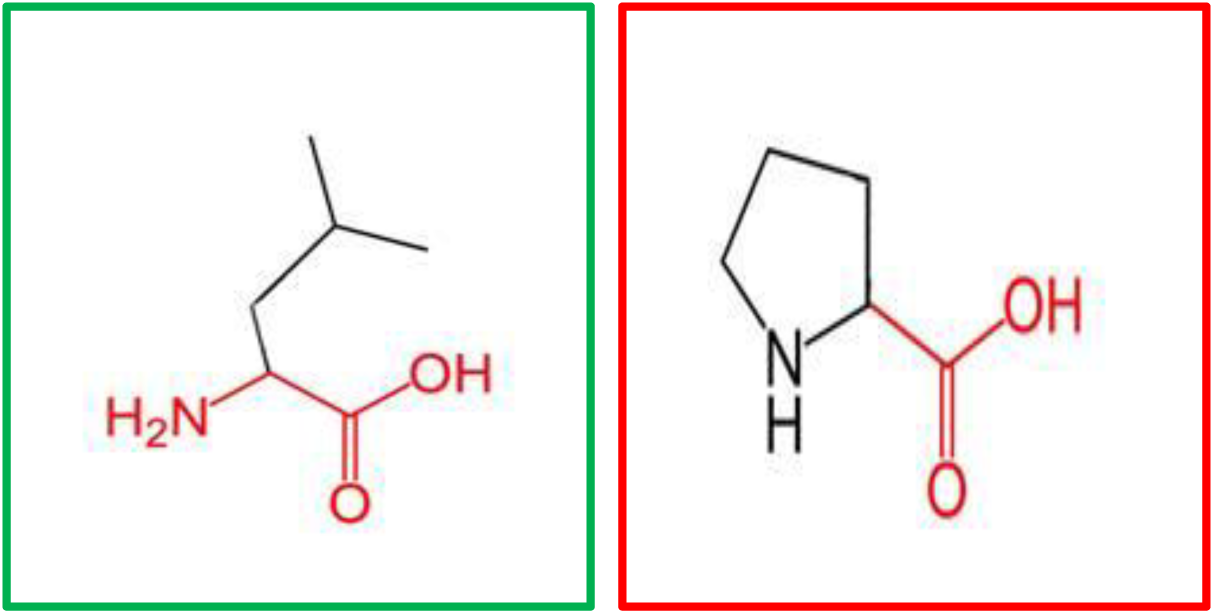
(L86P): change in the amino acid Leucine (green box) into Proline (red box) at position 86.

**Figure 4:**
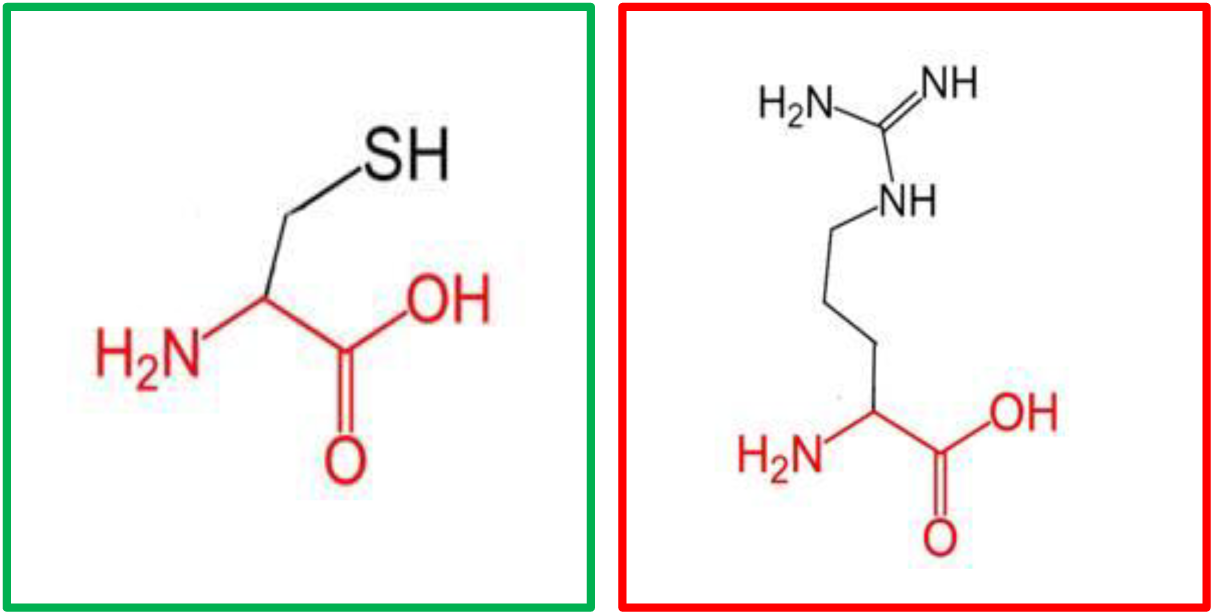
(C375R): change in the amino acid Cysteine (green box) into Arginine (red box) at position 375.

**Figure 5:**
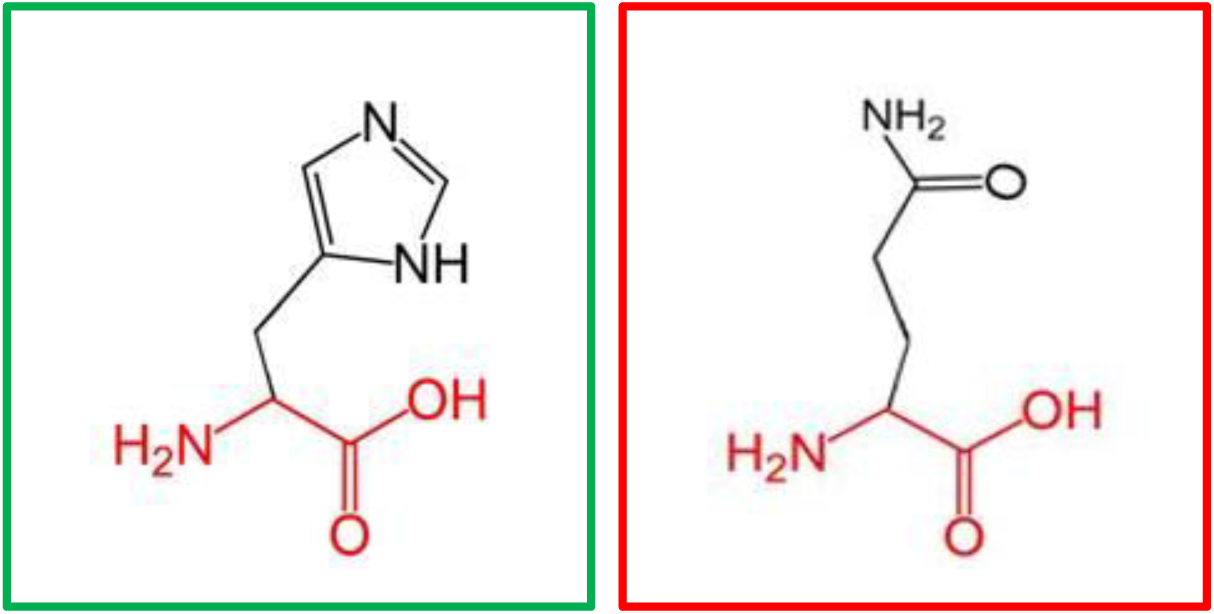
(H378Y): change in the amino acid Histidine (green box) into Tyrosine (red box) at position 378.

**Figure 6:**
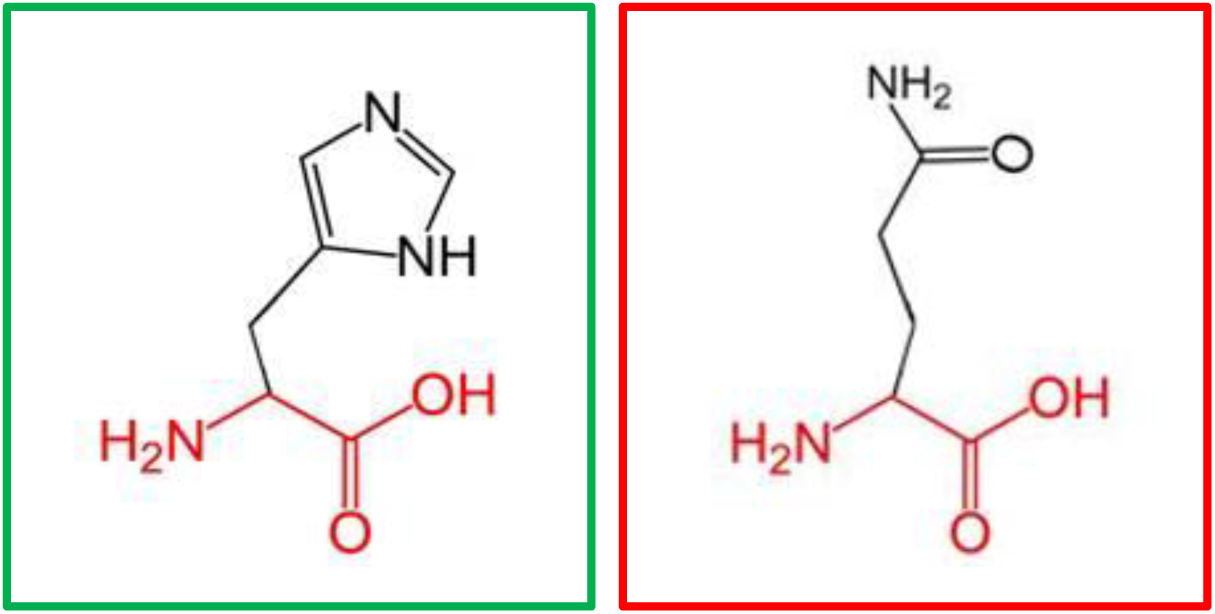
(H378Q): change in the amino acid Histidine (green box) into Glutamine (red box) at position 378.

**Figure 7:**
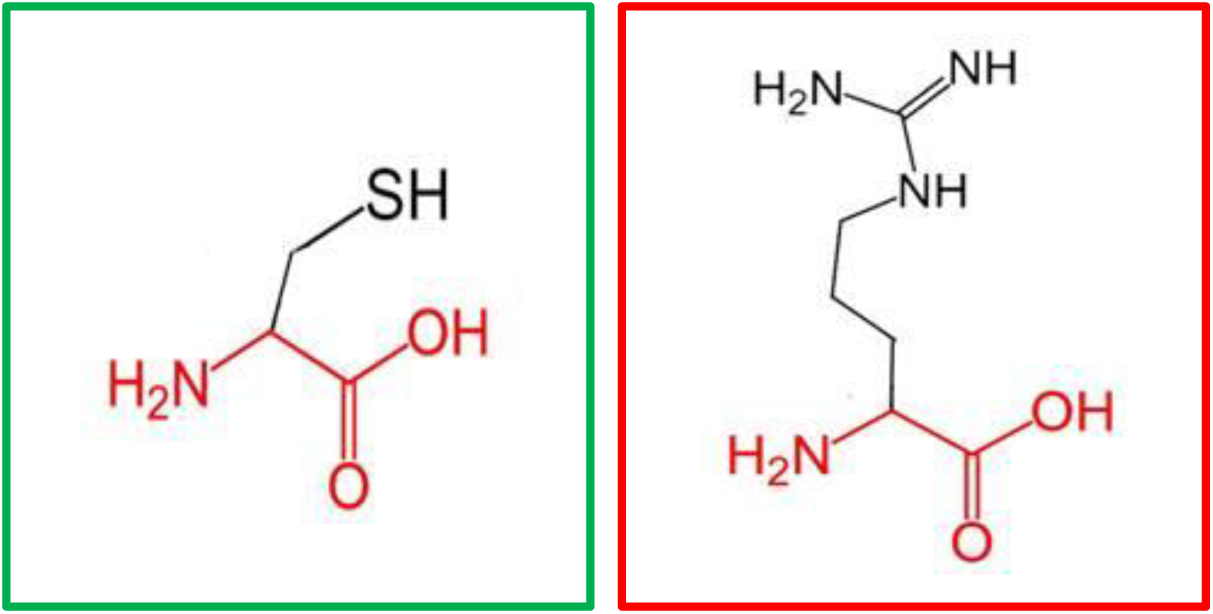
(C395R): change in the amino acid Cysteine (green box) into Arginine (red box) at position 395.

**Figure 8:**
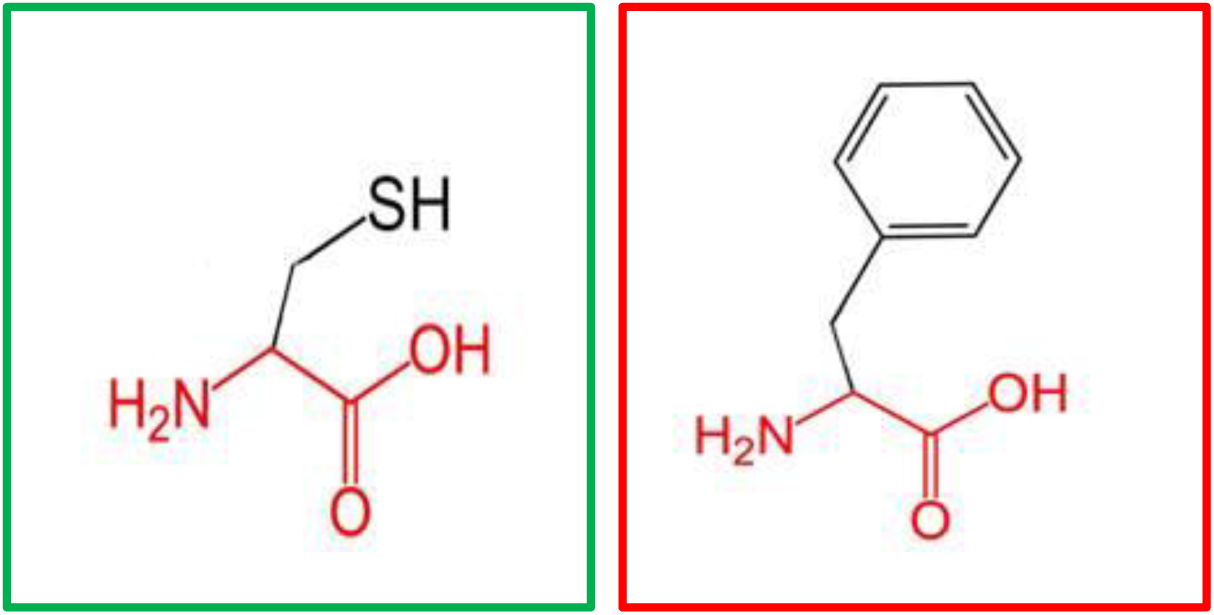
(C395F): change in the amino acid Cysteine (green box) into Phenylalanine (red box) at position 395.

**Figure 9:**
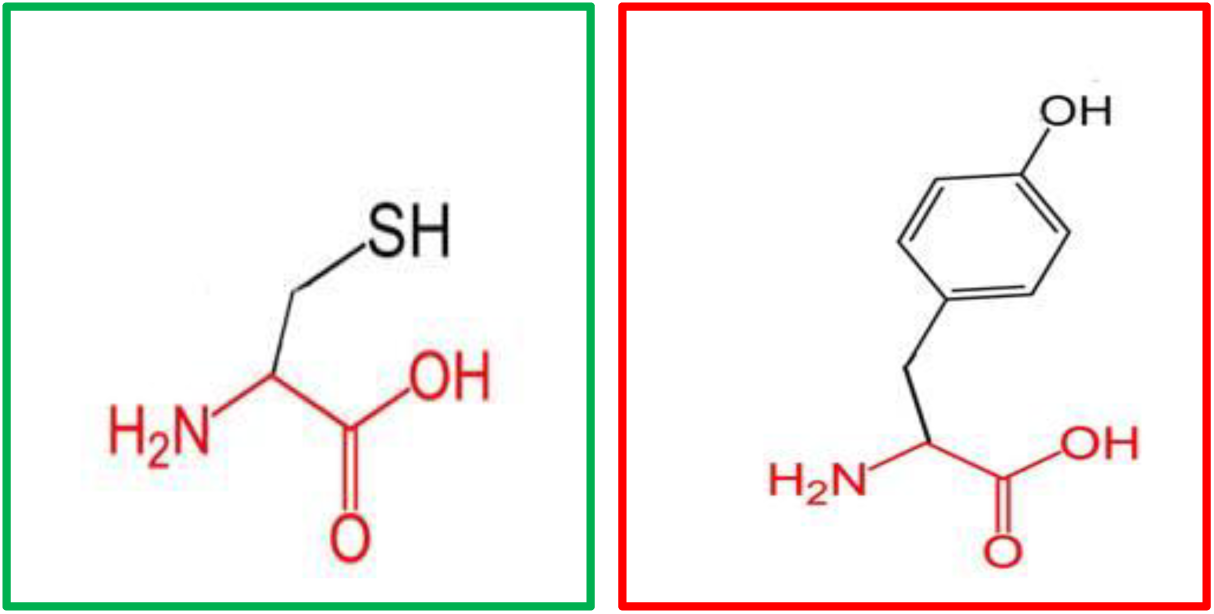
(C395Y): change in the amino acid Cysteine (green box) into Tyrosine (red box) at position 395.

**Figure 10:**
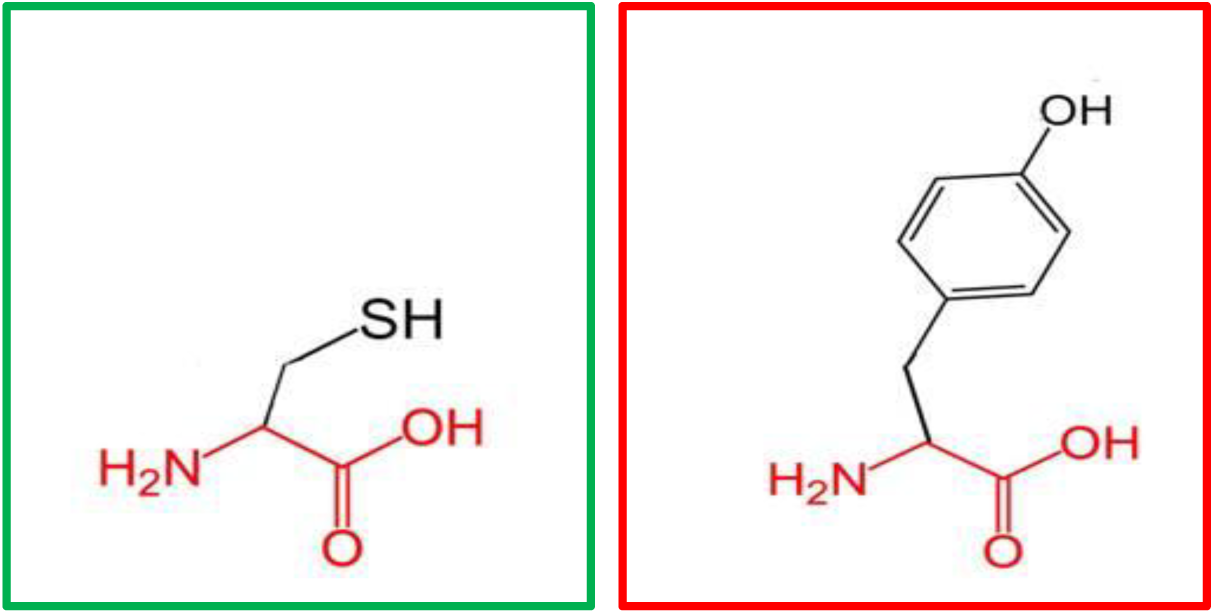
(C398Y): change in the amino acid Cysteine (green box) into Tyrosine (red box) at position 398.

**Figure 11:**
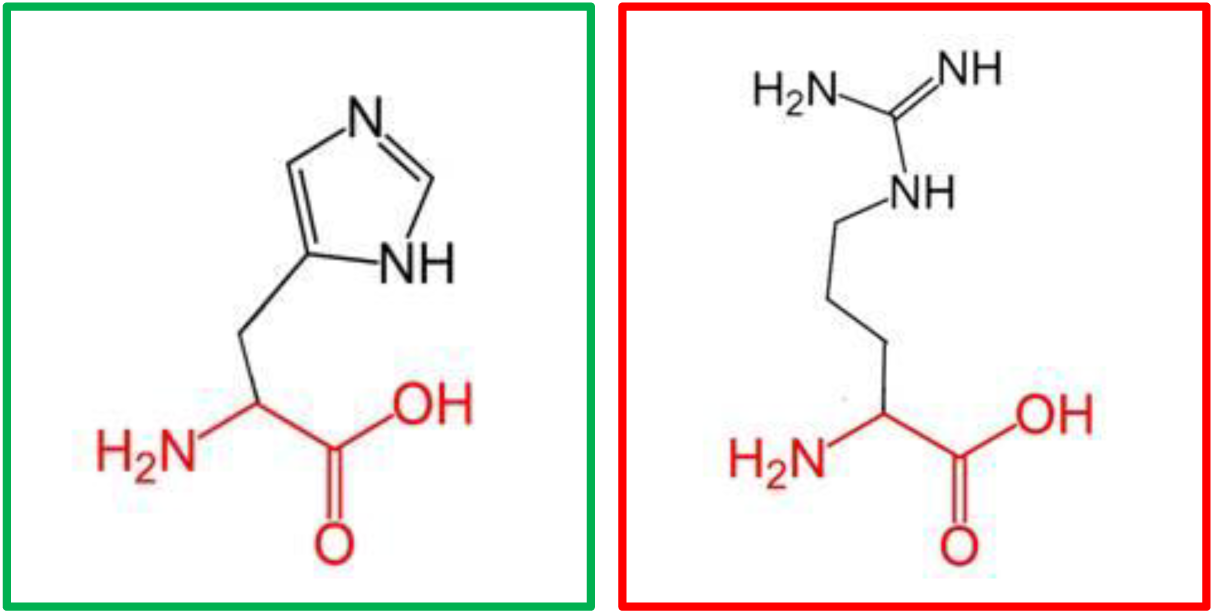
(H404R): change in the amino acid Histidine (green box) into Arginine (red box) at position 404.

**Figure 12:**
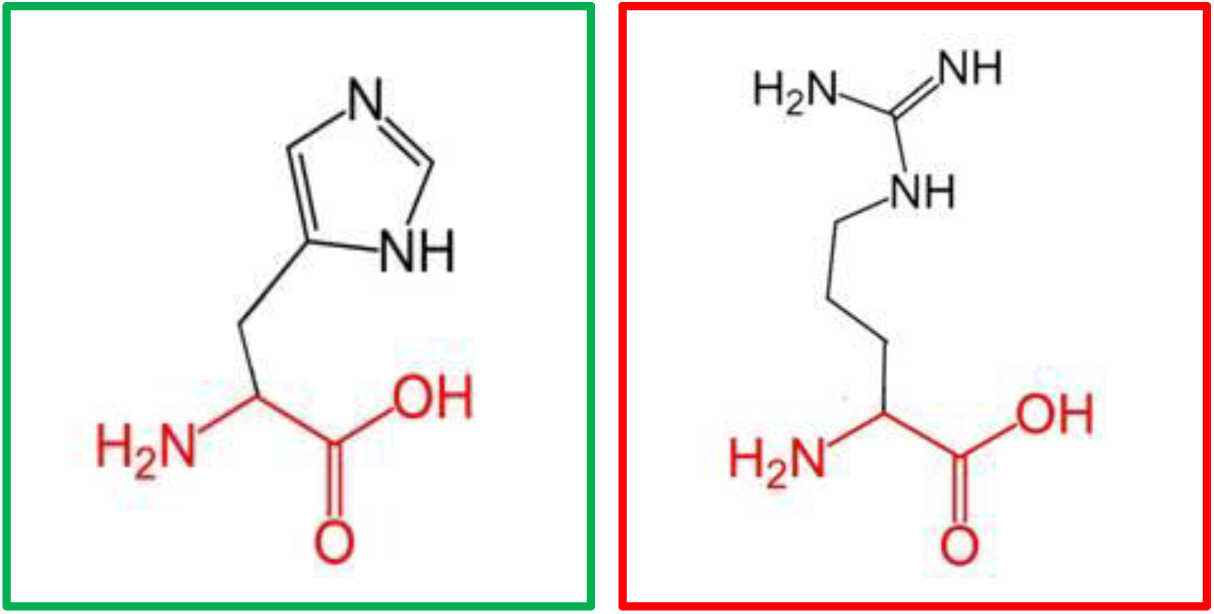
(H407R): change in the amino acid Histidine (green box) into Arginine (red box) at position 407.

**Figure 13:**
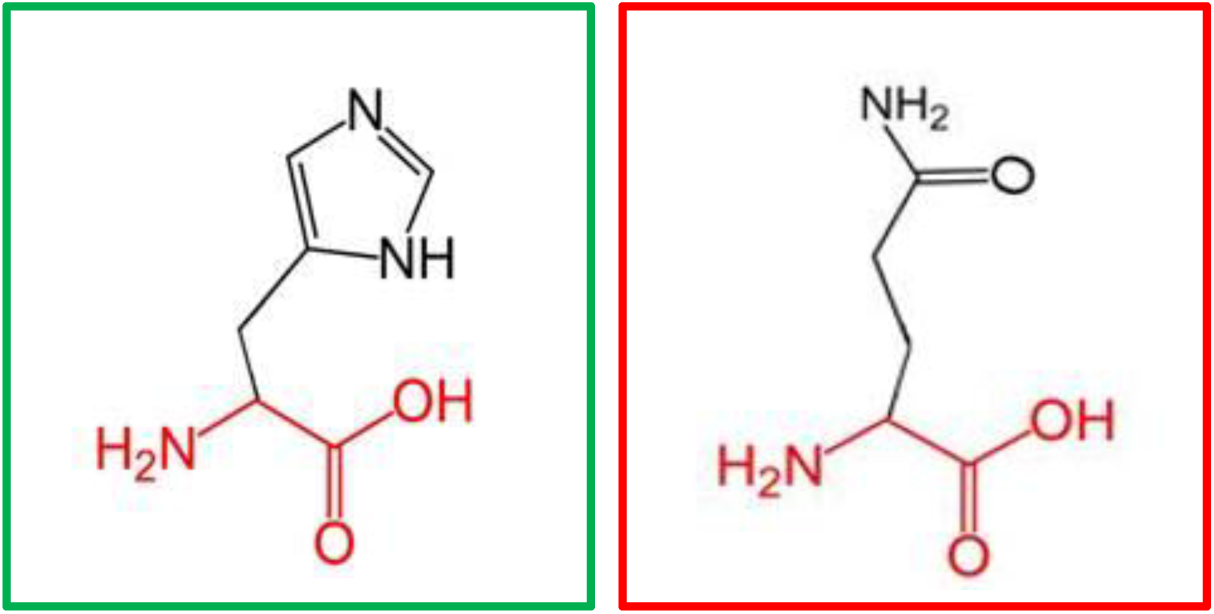
(H407Q): change in the amino acid Histidine (green box) into Glutamine (red box) at position 407.

**Figure 14:**
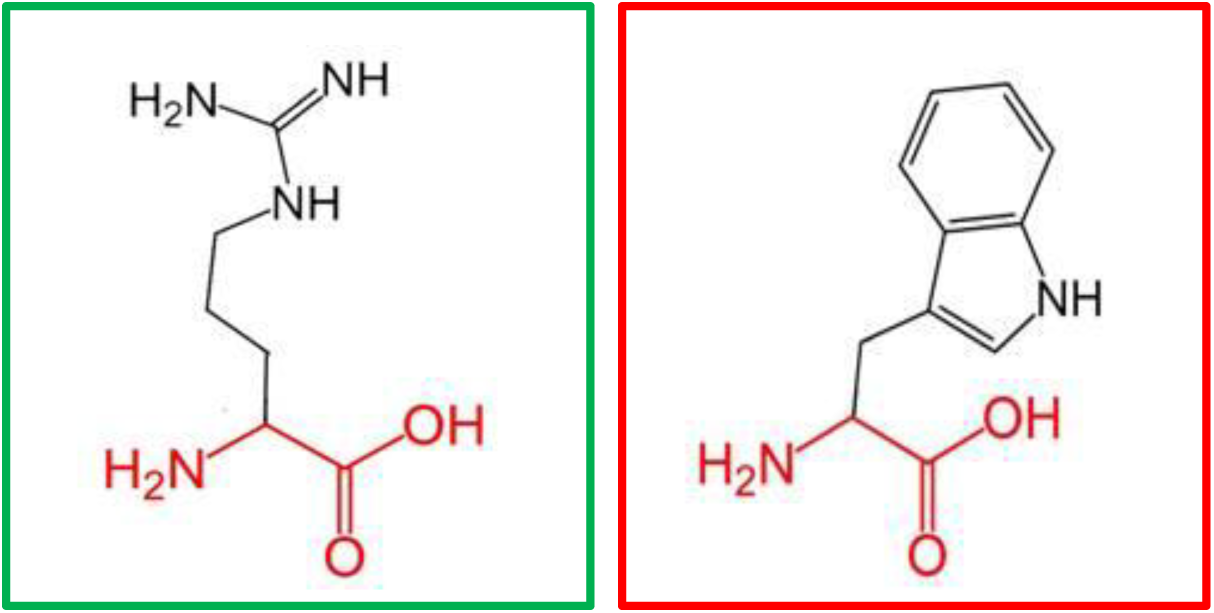
(R461W): change in the amino acid Arginine (green box) into Tryptophan (red box) at position 461.

**Figure 15:**
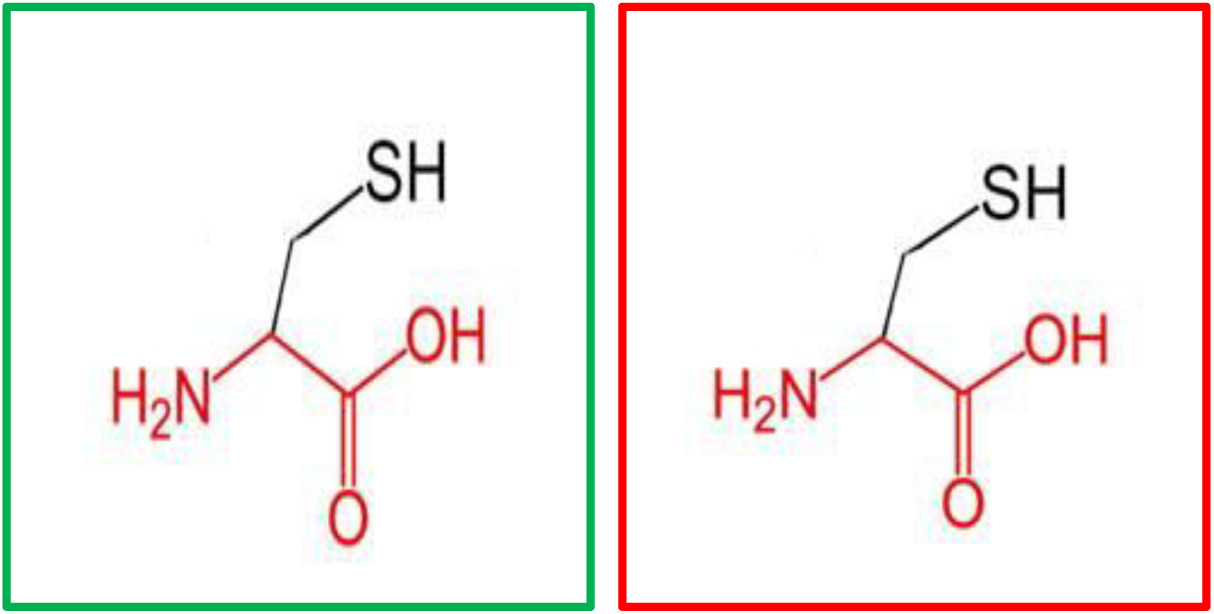
(F636C): change in the amino acid Phenylalanine (green box) into Cysteine (red box) at position 636.

**Figure 16:**
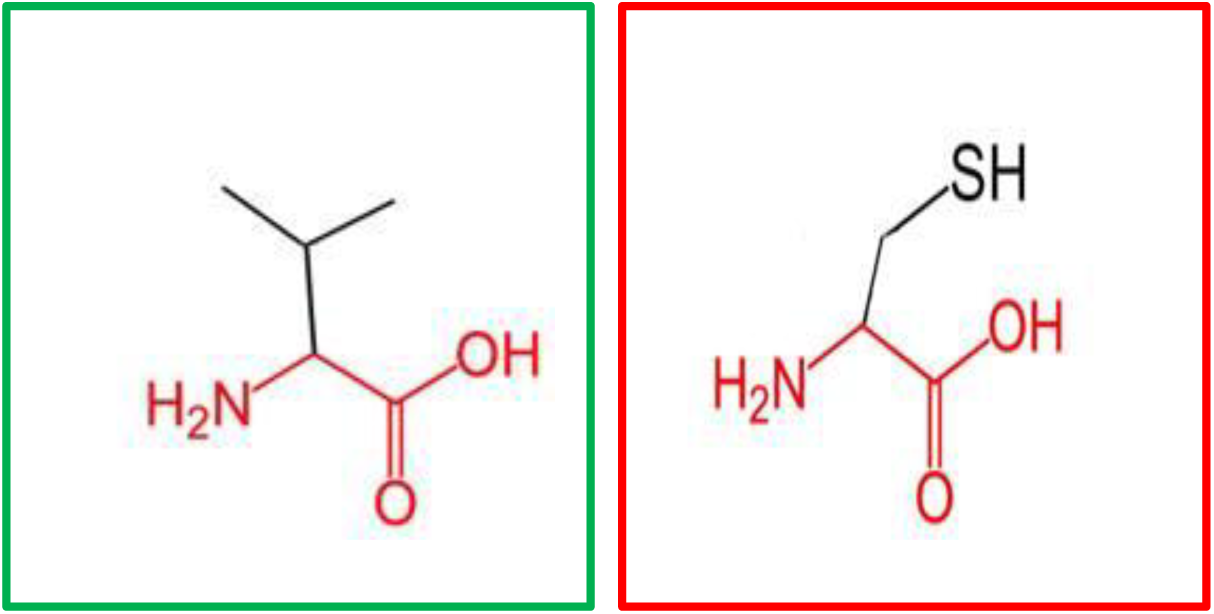
(V659F): change in the amino acid Valine (green box) into Phenylalanine (red box) at position 636.

**Figure 17:**
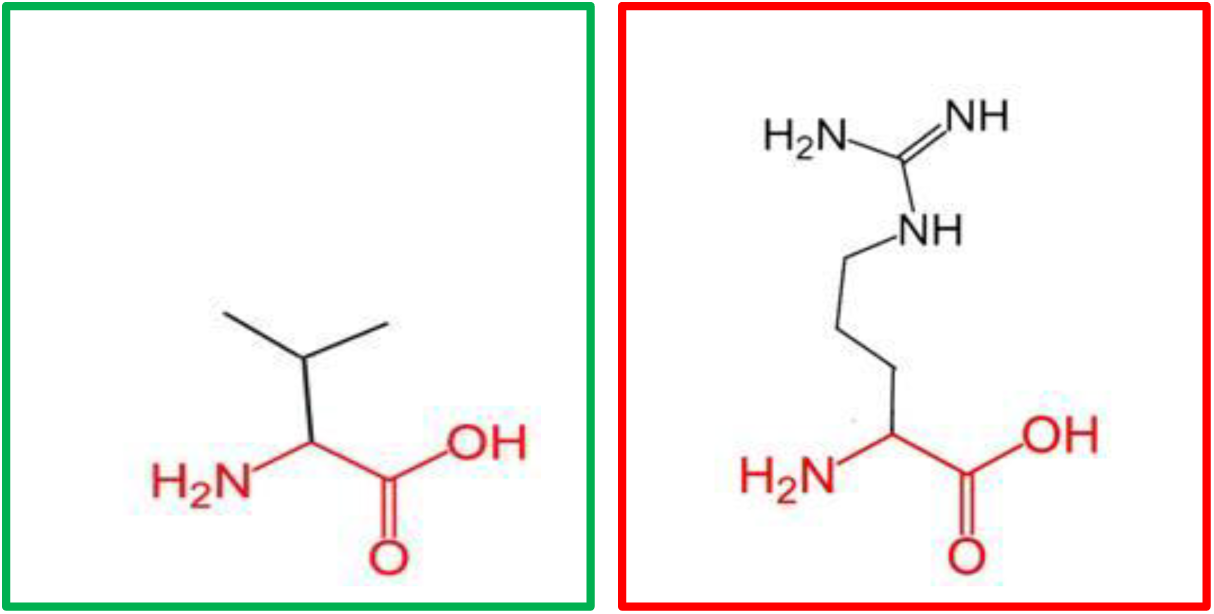
(G668R): change in the amino acid Glycine (green box) into Arginine (red box) at position 668.

**Figure 18:**
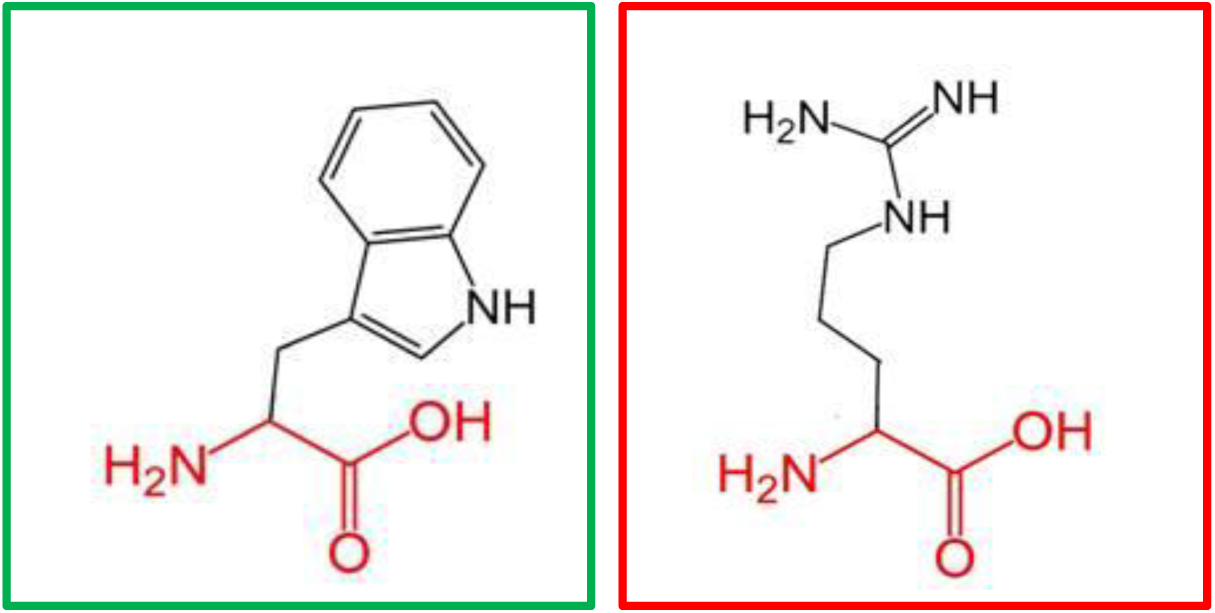
(W689R): change in the amino acid Tryptophan (green box) into Arginine (red box) at position 689.

**Figure 19:**
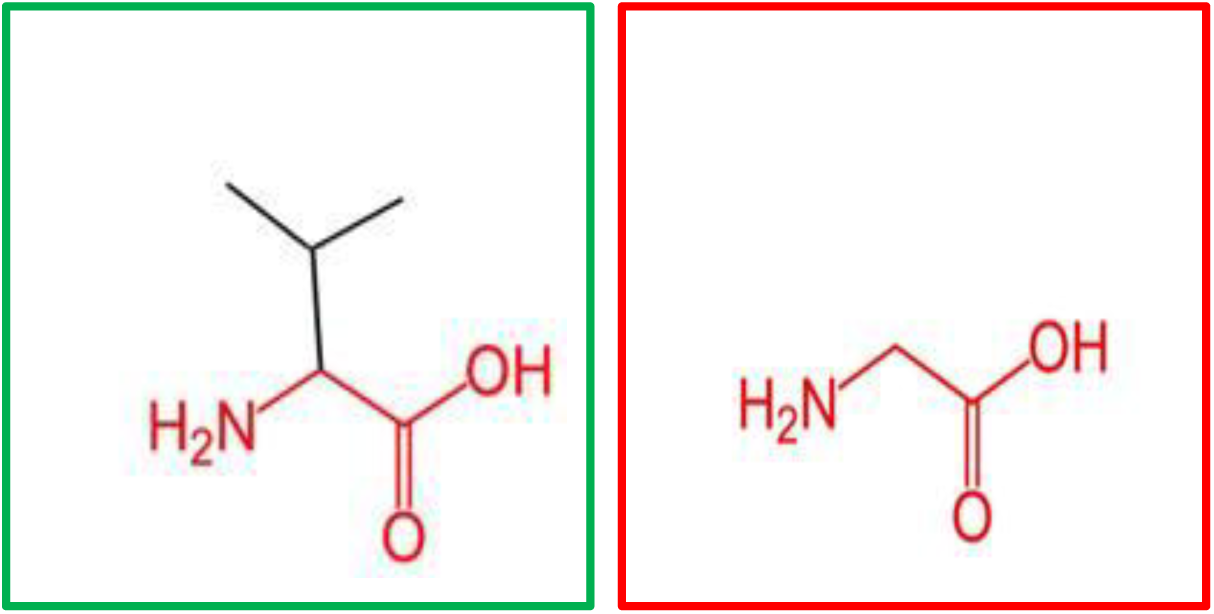
(V691G): change in the amino acid Valine (green box) into Glycine (red box) at position 691.

**Figure 20:**
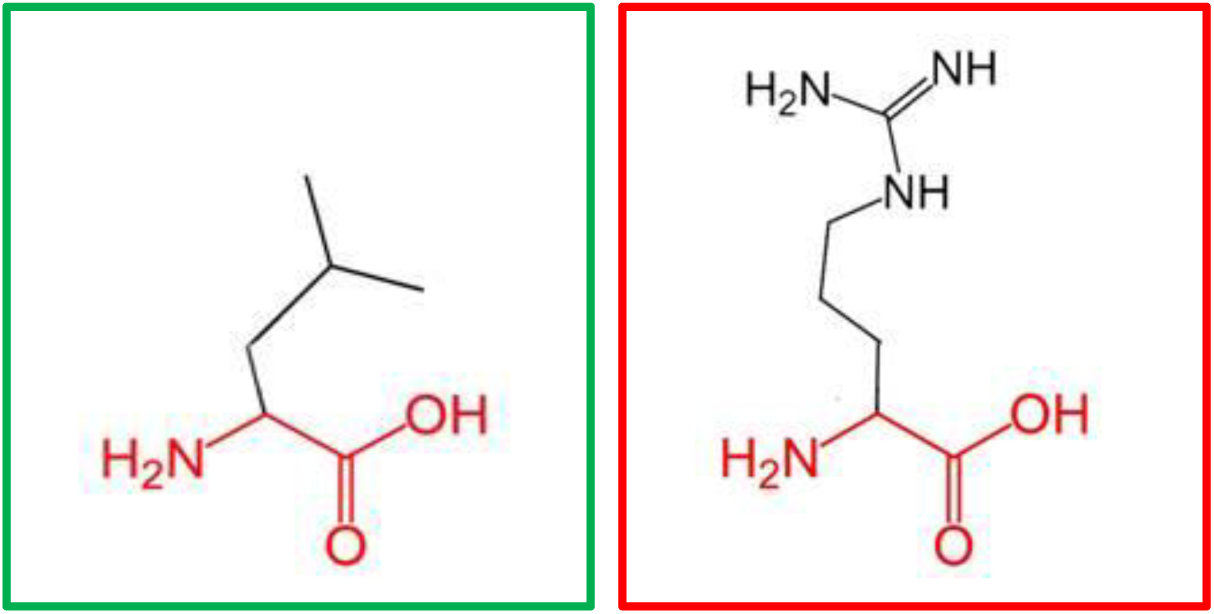
(L709R): change in the amino acid Leucine (green box) into Arginine (red box) at position 709.

**Figure 21:**
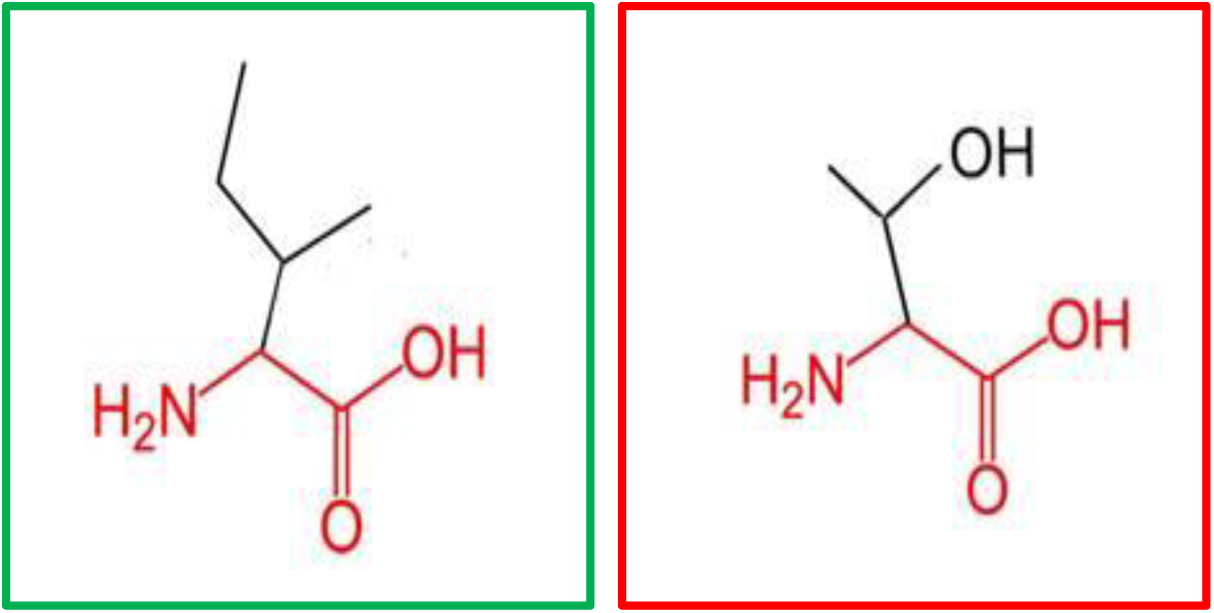
(I720T): change in the amino acid Isoleucine (green box) into Threonine (red box) at position 720.

**Figure 22:**
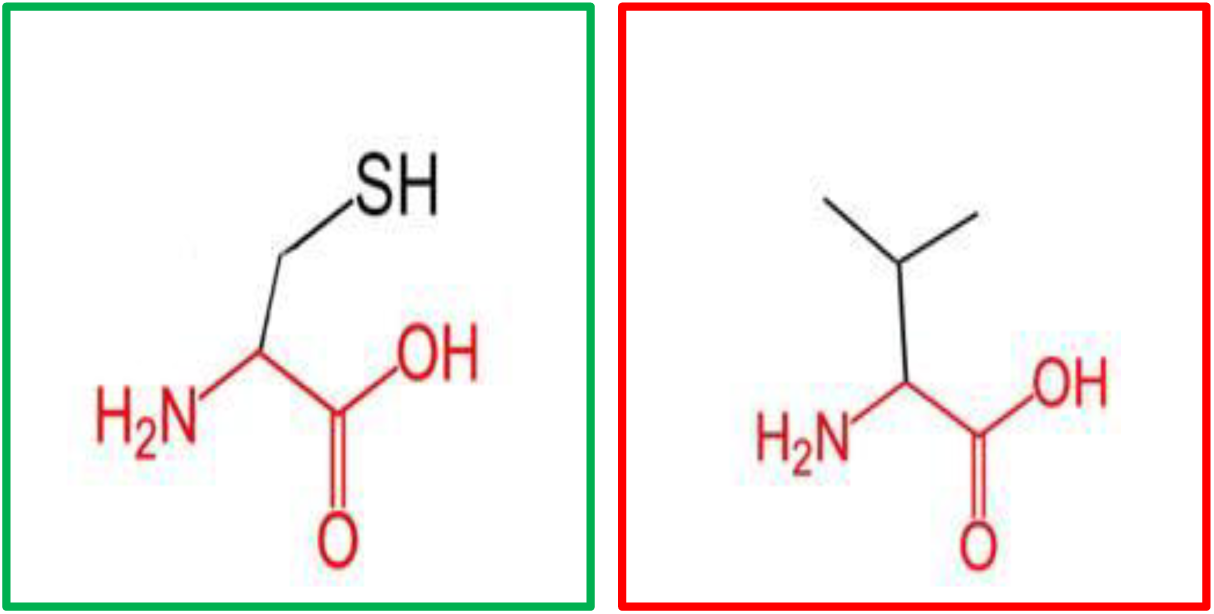
(F731V): change in the amino acid Phenylalanine (green box) into Valine (red box) at position 731.

**Figure 23:**
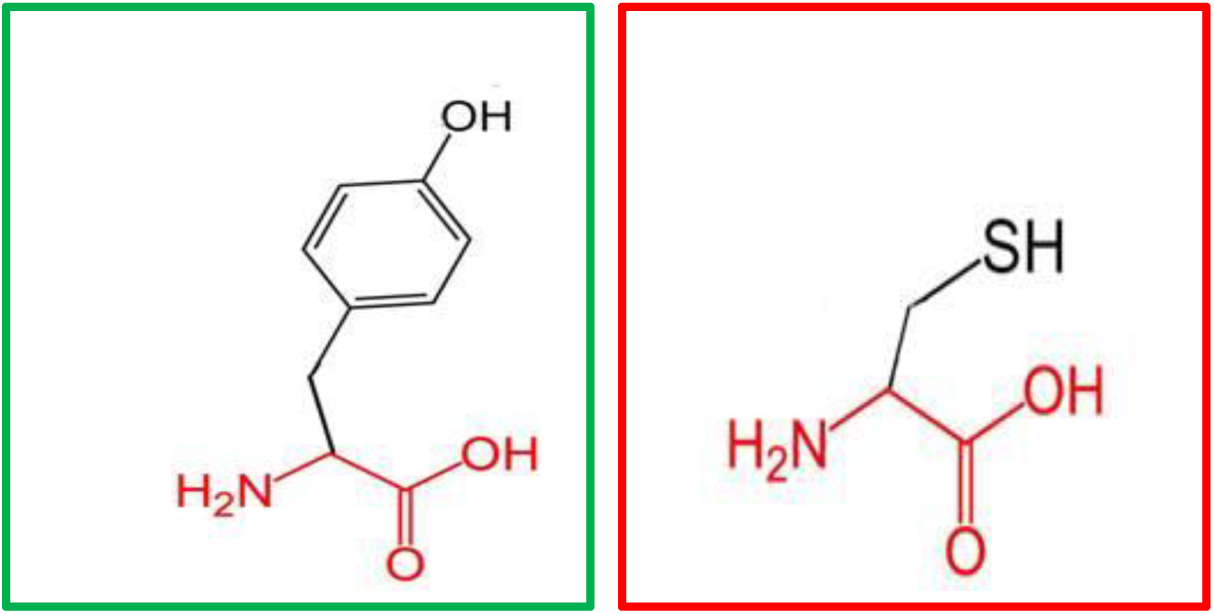
(Y741C): change in the amino acid Tyrosine (green box) into Cysteine (red box) at position 731.

**Figure 24:**
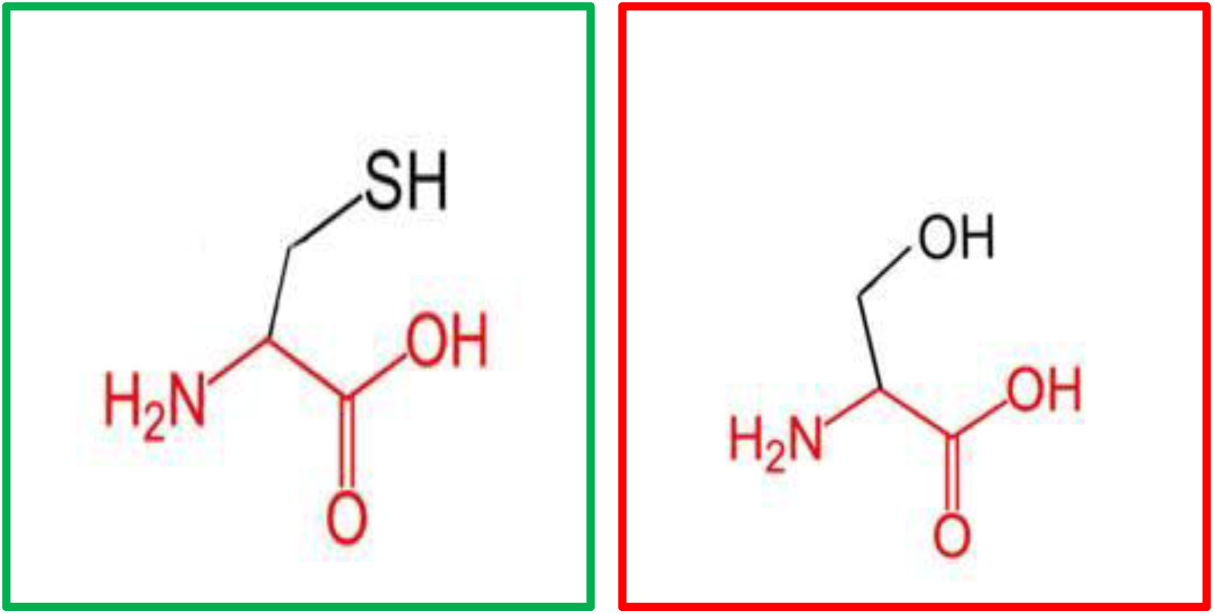
(F743S): change in the amino acid Phenylalanine (green box) into Serine (red box) at position 743.

**Figure 25:**
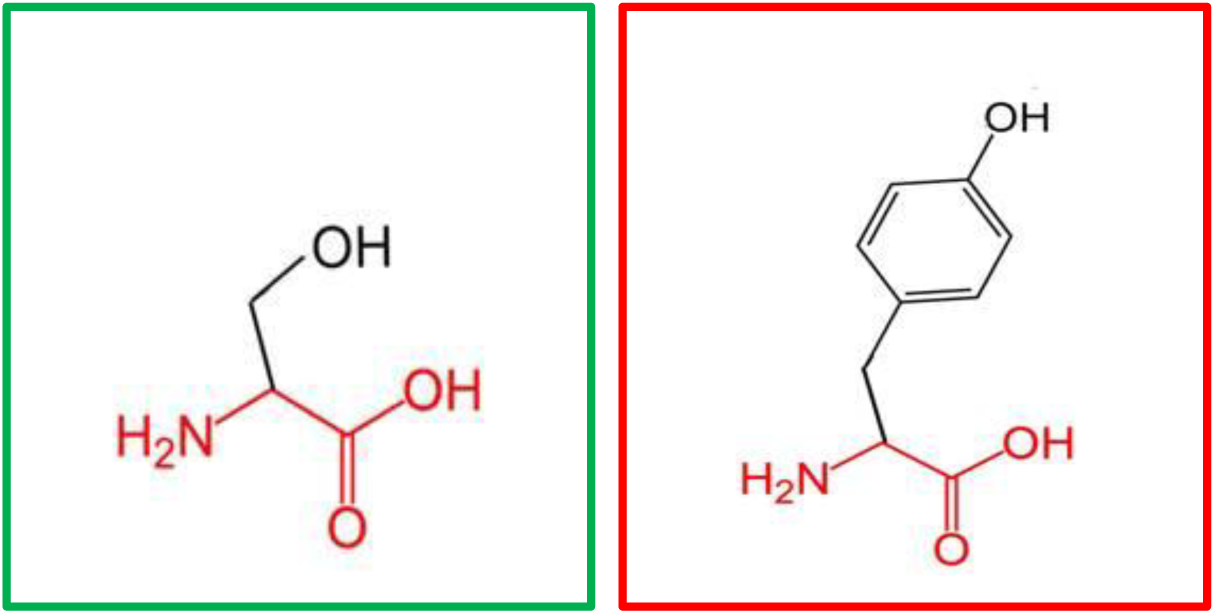
(S749Y): change in the amino acid Serine (green box) into Tyrosine (red box) at position 749.

GeneMANIA revealed that *MEFV* has many vital functions: chemokine production, inflammatory response, interleukin-1 beta production, interleukin-1 production, intracellular receptor signaling pathway, nucleotide-binding domain, leucine rich repeat containing receptor signaling pathway, positive regulation of cysteine-type endopeptidase activity, positive regulation of endopeptidase activity, positive regulation of peptidase activity, regulation of chemokine production, regulation of cysteine-type endopeptidase activity, regulation of endopeptidase activity, regulation of interleukin-1 beta production, regulation of interleukin-1 production, regulation of peptidase activity. The genes co-expressed with, share similar protein domain, or participate to achieve similar function were illustrated by GeneMANIA and shown in figure (26) Table (4 & 5).

**Figure 26:**
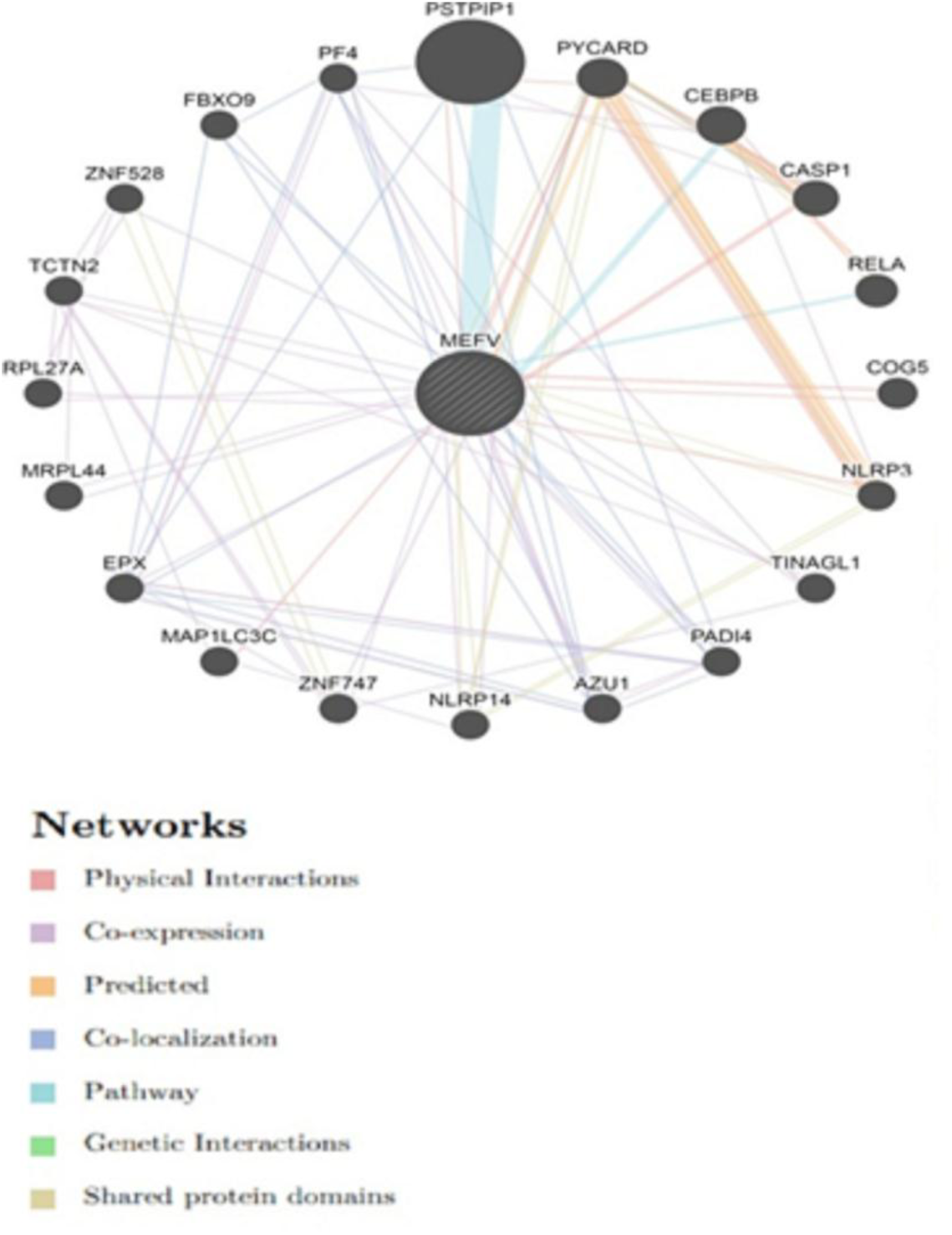
Interaction between *MEFV* and its related genes.

**Table (4):**
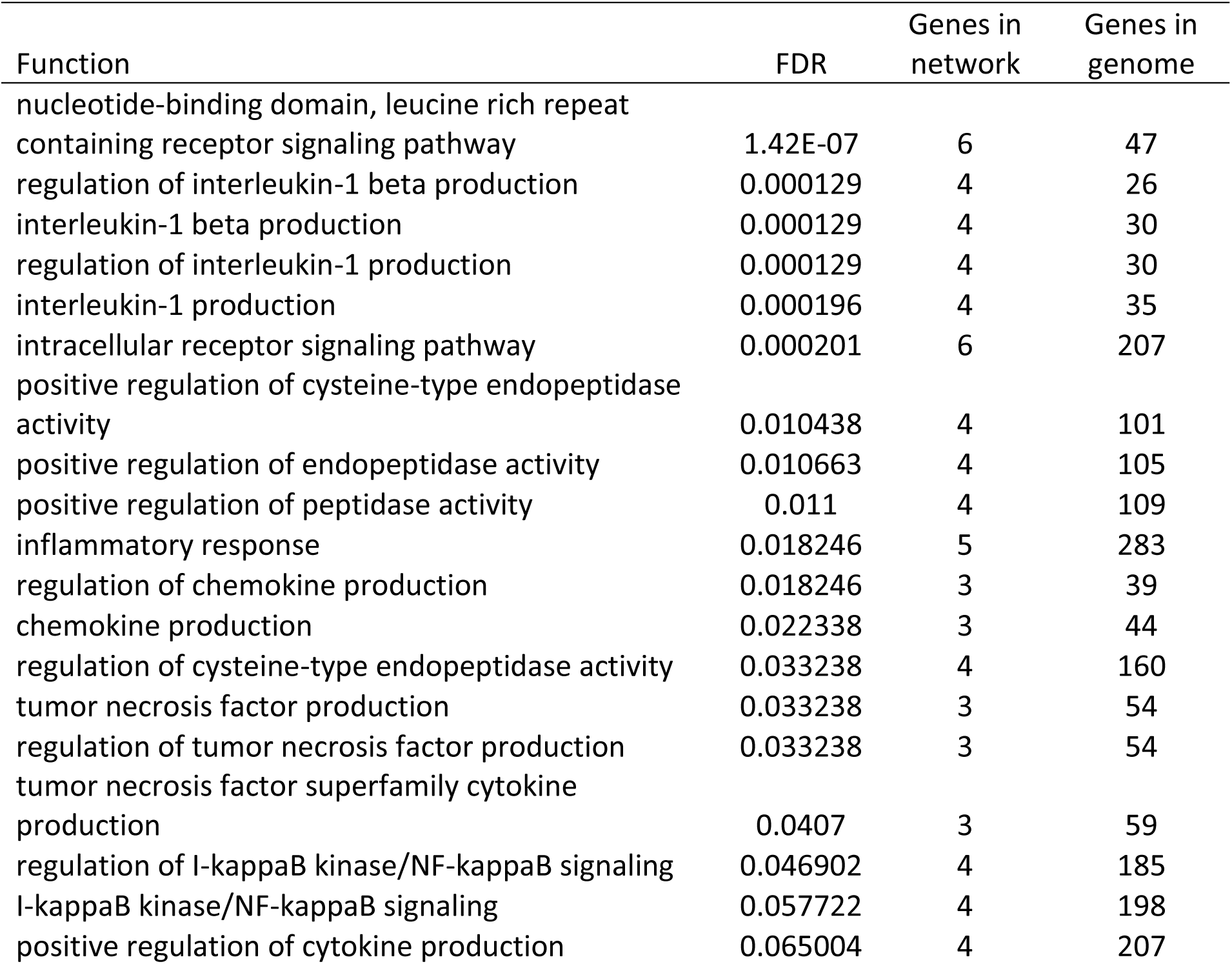
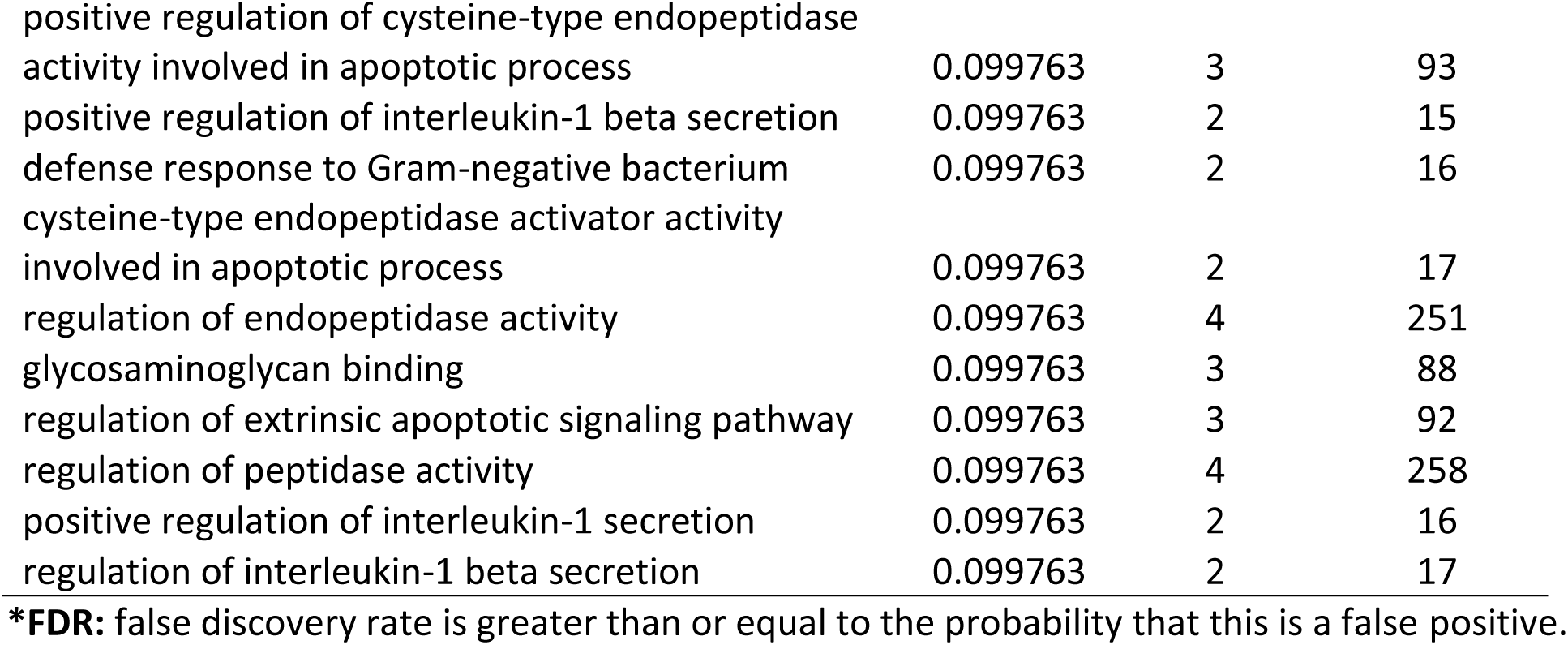
The *MEFV* gene functions and its appearance in network and genome:

**Table (5).**
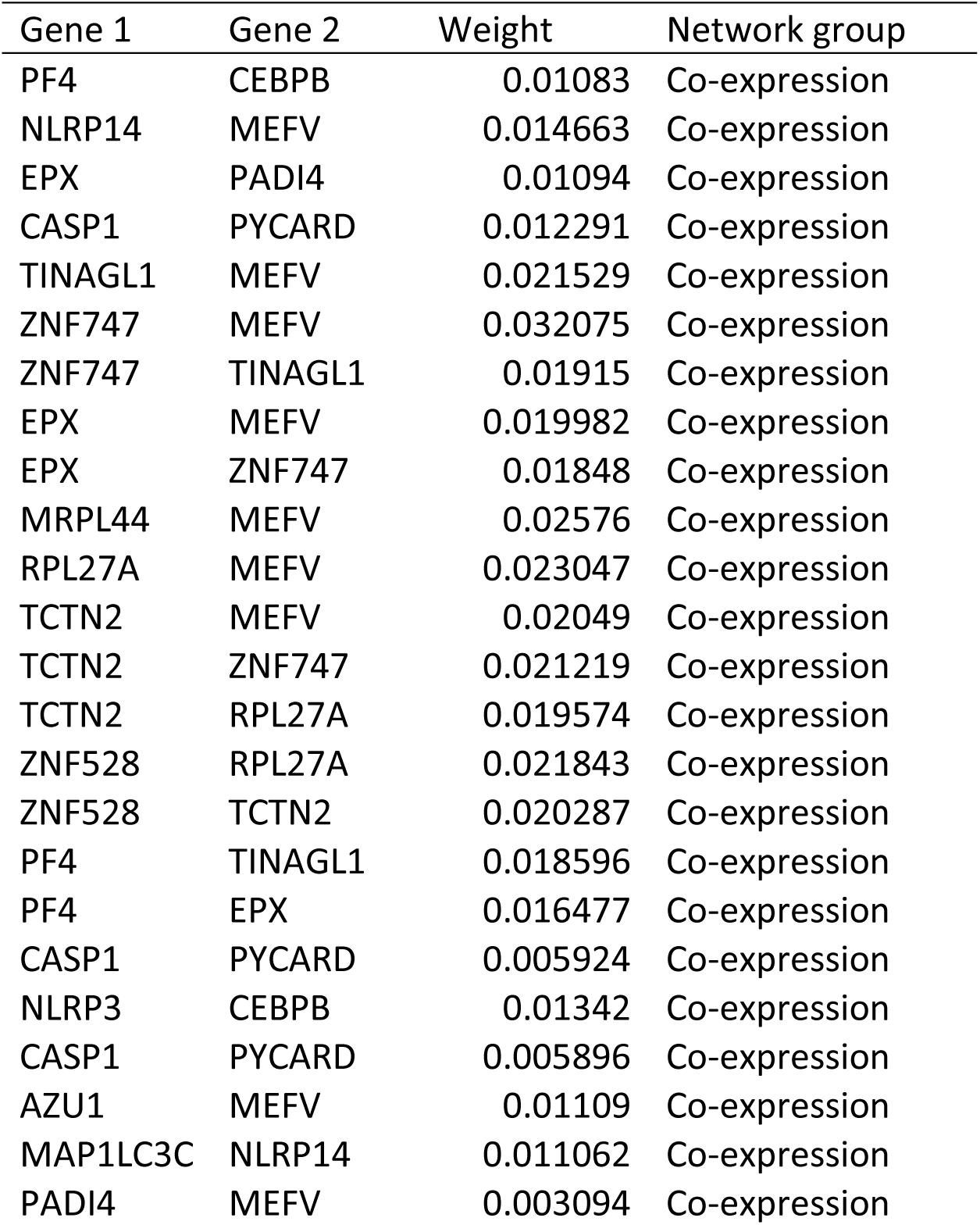
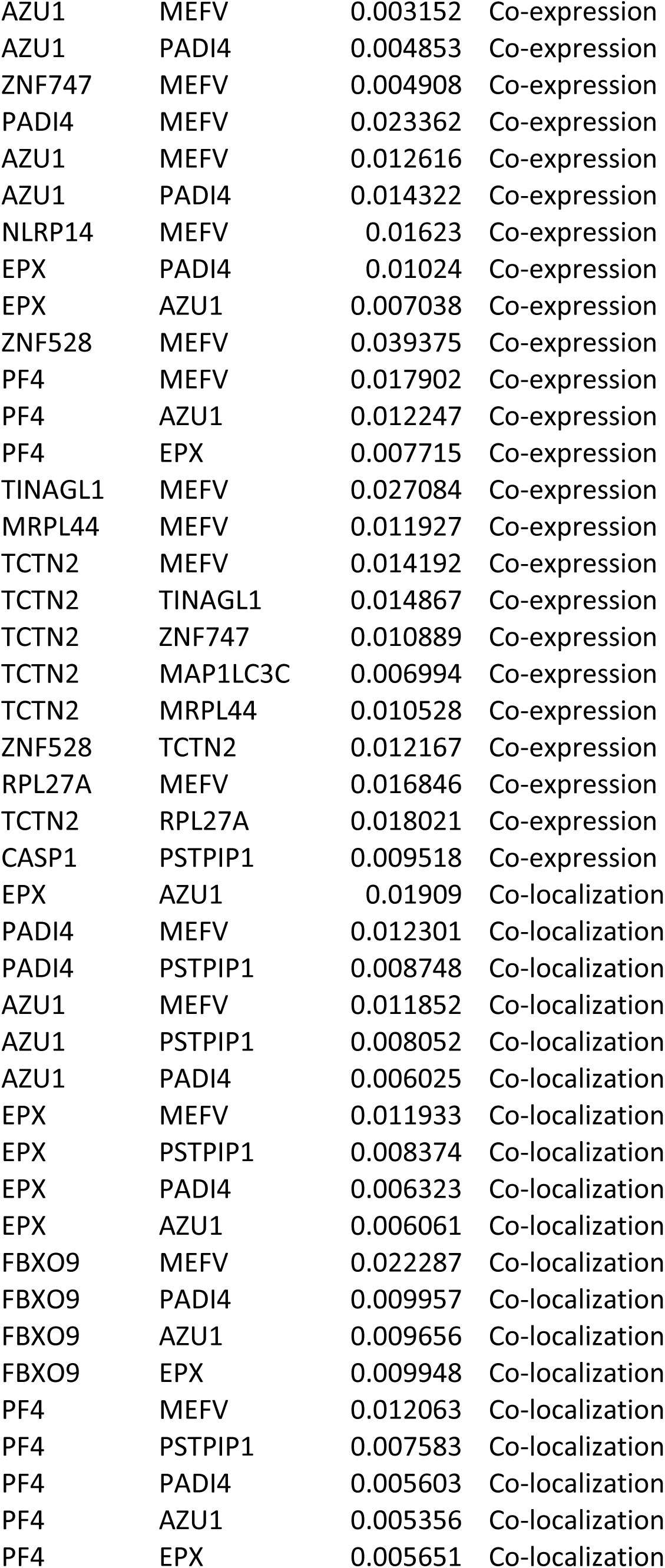
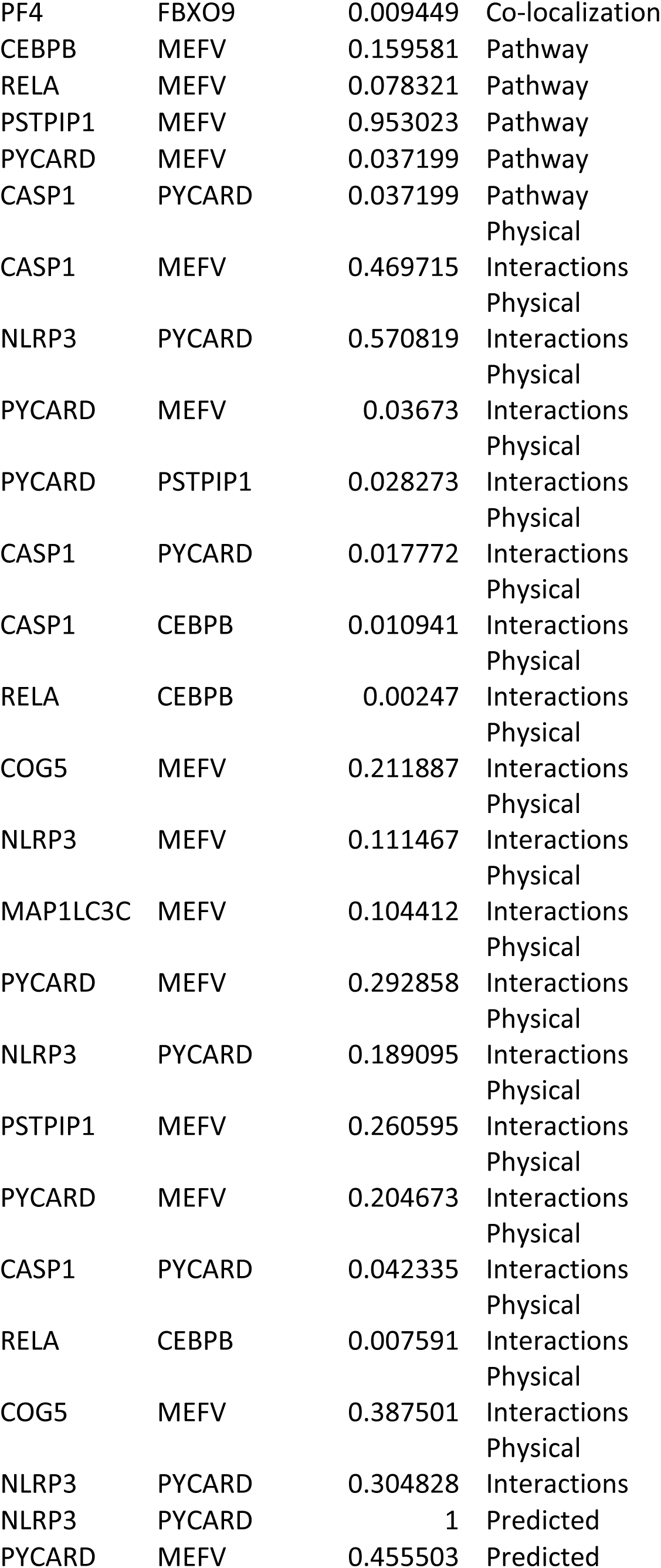
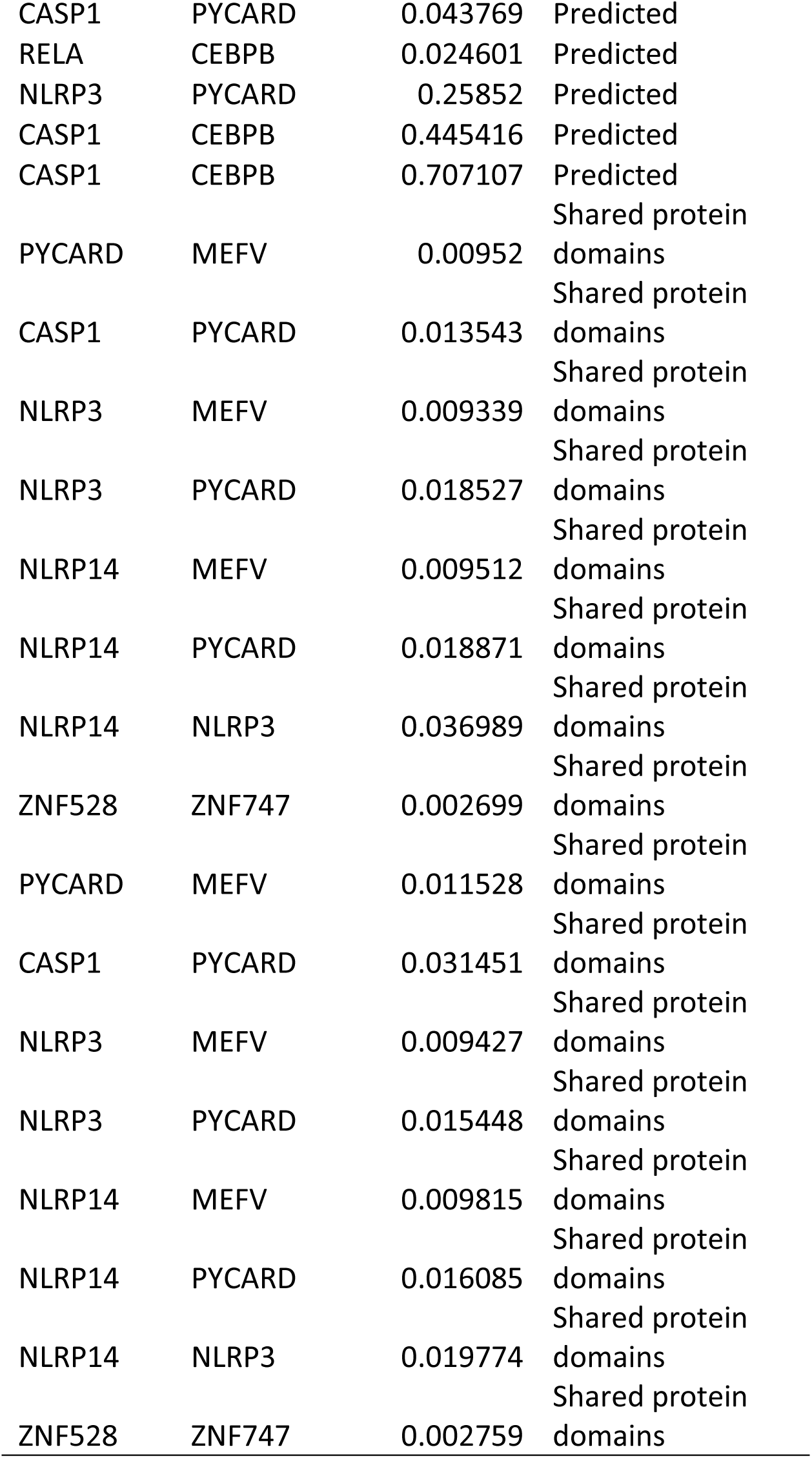
The gene co-expressed, share domain and Interaction with *MEFV* gene network:

In this study we also observed that, only one SNP (L86P) was found in conserve region, figure (3). The 10 amino acid sequences of *MEFV* gene were retrieved from UniProt (34). Sequence Alignments were done by BioEdit (v7.2.5). We also found that (H404R) pathogenic, which is matched with the result that we retrieved from dbSNPs/NCBI database. also we Retrieved all these SNPs as untested (V659F, L709R, F743S, S749Y) we found to be all damaging. Our study is the first in silico analysis of *MEFV* gene which was based on functional analysis while all previous studies (35, 36) based on frequency. This study revealed 23 Novel Pathological Mutations have a potential functional impact and may thus be used as diagnostic markers for Mediterranean basin populations.

## 4. Conclusion

In this work the influence of functional SNPs in the *MEFV* gene was investigated through various computational methods, determined that (S749Y, F743S, Y741C, F731V, I720T, L709R, V691G, W689R, G668R, V659F, F636C, R461W, H407Q,, H407R, H404R, C398Y, C395Y, C395F, C395R, H378Q, H378Y, C375R, L86P) are new SNPs have a potential functional impact and can thus be used as diagnostic markers. Constitute possible candidates for further genetic epidemiological studies with a special consideration of the large heterogeneity of *MEFV* SNPs among the different populations.

## Acknowledgment

The authors wish to acknowledgment the enthusiastic cooperation of Africa City of Technology - Sudan.

## Authors contributions

Abstract: Mujahed I. Mustafa

Introduction: Fatima A. Abdelrhman

Methodology: Mujahed I. Mustafa

Result & Discussion: Mujahed I. Mustafa

Conclusion: Soada Ahmed Osman

Writing the original draft: Mujahed I. Mustafa

